# SOLQC : Synthetic Oligo Library Quality Control Tool

**DOI:** 10.1101/840231

**Authors:** Omer Sabary, Yoav Orlev, Roy Shafir, Leon Anavy, Eitan Yaakobi, Zohar Yakhini

## Abstract

**Motivation:** Recent years have seen a growing number and a broadening scope of studies using synthetic oligo libraries for a range of applications in synthetic biology. As experiments are growing by numbers and complexity, analysis tools can facilitate quality control and help in assessment and inference.

**Results:** We present a novel analysis tool, called **SOLQC**, which enables fast and comprehensive analysis of synthetic oligo libraries, based on NGS analysis performed by the user. SOLQC provides statistical information such as the distribution of variant representation, different error rates and their dependence on sequence or library properties. SOLQC produces graphical descriptions of the analysis results. The results are reported in a flexible report format. We demonstrate SOLQC by analyzing literature libraries. We also discuss the potential benefits and relevance of the different components of the analysis.

**Availability:** https://app.gitbook.com/@yoav-orlev/s/solqc/

## 1 Introduction

DNA synthesis technology has greatly developed over recent years and is holding a promise to enable a leap in using natural systems for various applications. For example, synthetic DNA is used for making protein therapeutics and drugs. Another application is in genome editing, wherein optimizing CRISPR-Cas9 systems and reagents is enabled by using libraries of synthetic DNA oligonucleotides. Synthetic DNA is particularly useful for screening of large guide-RNA libraries to optimize CRISPR-Cas9 based systems [19]. The use of synthetic DNA also enables the optimization of crops for efficient biofuel production [24]. In particular, synthetic DNA can be used to perform codon-optimization, directed evolution, enzyme libraries screens, and incorporation of non-natural amino acids to improve novel enzymatic activities in the biofuel industry [8]. Last but not least, synthetic DNA is also an attractive alternative for data storage media, see e.g. [20] and more details in section 1.1. With an information density orders of magnitude better than that of magnetic media and due to its highly robust chemical properties DNA can potentially efficiently store data for centuries. This progress in the use of synthetic DNA, as well as its potential to drive future applications, drives work focused on the optimization of manufacturing processes and of design cycles. A key to such work is monitoring the quality of synthetic DNA throughout the process, including at the hands of the end users.

Synthetic DNA libraries consisting of thousands of DNA sequences, often referred to as **variants**, have become a common tool in molecular biology. They allow for systematic, unbiased investigations for discovery biology, directed evolution for protein engineering, and in vitro molecular optimization to generate mutant proteins with novel properties. Sharon et al. used an oligonucleotide library (OL) to infer gene regulatory logic [23], Levy et al. used an OL to discover a bacterial insulation mechanism [18], and most recently Kotler et al. found links between differential functional impact to mutations in p53 using OLs [16].

The process of using OLs in such studies usually starts with a design file containing the DNA variants, which will be synthesized as millions of physical oligonucleotides (oligos). These oligos will typically be sequenced in one or more steps of the experimental process, producing results in the form of NGS output files (typically fastq). We refer to each sequenced synthesized strand as a read. To validate and/or optimize the results of the different steps in an OL based study, it is important to quality control all components to make sure that the results stem from the biology and not from noise, from confounding interference of from other biases related to synthesis and to sequencing.

### 1.1 DNA Storage Systems

The recent progress in synthesis and sequencing technologies has paved the way for the development of a non-volatile data storage technology based upon DNA molecules. A DNA storage system consists of three important components. The first is DNA synthesis. The stage at which the strands or DNA molecules that encode the data are produced (those strands called input strands, or variants). In order to produce strands with acceptable error rates, in a high throughput manner, the length of the strands is typically limited to no more than 250 nucleotides [2]. The second part is a storage container with compartments. This container stores the DNA strands. No order is assumed for this stage. Finally, sequencing is performed to read back a representation of the strands (the output of this stage consists of strands called output strands or sequencing reads). A decoding process transforms the sequencing output back to digital data. The encoding and decoding stages are two processes, external to the storage system, that convert the user’s binary data into strands of DNA in such a way that, even in the presence of errors, it will be possible to revert back and reconstruct the original binary data of the user.

One of the first experiments to store information in DNA was conducted by Clellan et al. in 1999, where they coded and recovered a message consisting of 23 characters [7]. Shortly after this, in 2000, Leier et al. have managed to successfully store three sequences of nine bits each [17]. A more significant progress, in terms of the amount of data stored successfully, was reported by Gibson et al. in 2010, demonstrating in-vivo storage of 1,280 characters in a bacterial genome [10]. The first large scale demonstrations of the potential of in vitro DNA storage were reported by Church et al. who recovered 643 KB of data [6] and by Goldman et al. who accomplished the same task for a 739 KB message [11]. Both of these pioneering groups did not recover the entire message successfully as no error correcting codes were used. Later, in [12], Grass et al. stored and recovered successfully 81 KB message, in an encapsulated media, and Bornholt et al. demonstrated storing a 42 KB message [4]. A significant improvement in volume was reported in [3] by Blawat et al. who successfully stored 22 MB of data. Recently, Erlich and Zielinski managed to store 2.11 MB of data with high storage density [9]. The largest volume of stored data is reported by Organick et al. in [20]. Organick et al. describe the encoding and decoding of 200 MB of data, an order of magnitude more data than previously reported. Yazdi et al. [28] developed a method that offers both random access and rewritable storage. Most recently, Anavy et al. [1] described how more data can be stored for less synthesis cycles. Their approach uses composite DNA letters. A similar approach, on a smaller scale, was reported in [5].

### 1.2 Synthetic Oligo Library (OL) Errors

The processes of synthesizing, storing, sequencing and handling oligonucleotides are all error prone. Each step in the process can independently introduce a significant number of errors:

1. Both the synthesis process and the sequencing process can introduce deletions, insertions, and substitution errors on each of the reads and/or synthesized strands.
2. Current synthesis methods can not generate one copy for each design variant. They all generate thousands to millions of non perfect copies. Each of these copies has a different distribution of errors. Moreover, there might be some variants with a significantly larger number of copies, while some variants may be not be represented at all. In other words, the representation and the error profile of variants in the library is not uniform.
3. The use of DNA for storage or of OLs for other applications typically involves PCR amplification of the strands in the DNA pool [13]. PCR is known to have a preference for some sequences over others, which may further distort the distribution of the number of copies of individual sequences and their error profiles [21, 22].

Most of the research on characterizing errors in synthetic DNA libraries has been done in the context of individual studies using synthetic DNA. Tian et al. showed in [26] that the rate of deletion is 1/100 per position, insertion is 1/400 per position, and the rate for substitution is 1/400. Later, Kosuri and Church [15] noted that column-based oligo synthesis has total error rate of approximately 1/200 or less for oligos of 200 bases, where the most dominant error is a single base deletion. In addition, they showed that high GC content, at more than 50% of the bases in the strand being G or C, can inhibit the assembly and lead to lost data. They also pointed-out that in OL synthesis, a synthesis method based on DNA microarrays, the error rates are usually higher than those for column-based synthesis. Recently, in [13], Heckel, Mikutis, and Grass, studied the errors in a DNA storage channel based upon three different data sets from the experiments in [9, 11, 13]. In their work they studied the deletion/insertion/substitution rate and how it is affected by filtering reads with incorrect length (compared to the designed length). In particular, when they considered only reads with the correct length, they showed, as expected, that the deletion rate has been significantly decreased in all of the data sets. They also investigated the conditional error probability for substitutions and found out that in [9] the most dominant substitution error was from G to T (20%), and in the rest of the experiments, the most dominant substitution error was from C to G (about 30-40%). They also examined the effect of the number of PCR cycles on the coverage depth, which is the distribution of the number of reads per each of the variants. They concluded that, since the efficiency of the PCR amplification on each of the strands is different, a larger number of PCR amplification cycles leads to a higher differences in the coverage depth distribution of the variants. Organick et al. also characterized the errors in their experiment [20]. First, they found that substitution was the most frequent error in the library, then deletion, and lastly insertion. Furthermore, they found that while deletions showed almost equal rates for all of the four bases, insertions were mostly associated with base G, and substitutions were mostly associated with base T. Lastly, they also examined the read error rates per position. It should be noted that substitution errors are most likely associated with sequencing and not with synthesis.

### 1.3 This Work

In this paper we describe SOLQC, a software tool that supports the statistical analysis and quality control of OLs. The tool is designed to enable and to facilitate individual labs obtaining information about DNA libraries and performing error analysis before or during experiments. We describe our methods and demonstrate the results of analyzing several libraries from the literature. The dissemination reflects only the authors’ view and the EU Commission is not responsible for any use that may be made of the information it contains.

## 2 Materials and Methods

### 2.1 SOLQC Tool

In this section we present our software tool, called **SOLQC - Synthetic Oligo Library Quality Control**. This quality control tool generates a customized report which consists of several statistics and plots for a given input synthetic library. Detailed instructions to use the tool are given in Section 6.

The input to the SOLQC tool is the result of a sequencing reaction run on the library. It consists of the design variants and of all the sequenced reads. The input to the tool is provided using the following three files.

1. Design file: This file consists of the design variants that were synthesized and it has to be in a csv format. The tool also supports an IUPAC description [14] of the design.
2. NGS results file: This file is in fastq format and contains the NGS results.
3. Config file: Auxiliary file which consists of other details on the design variants such as information on the barcode etc.

The SOLQC tool is operated in the following order.

1. **Preprocessing**: The reads can be filtered such that only valid reads will be processed by the tool. The selection of valid reads can be configured by the user according to the sequence barcode and its length.
2. **Matching**: Each read is matched to its corresponding variant. The matching step can be done by different strategies as follows.

- Barcode matching: If the library has a barcode assigned to each variant, the barcode will be used in order to match each read with a tunable tolerance in errors for the matching.
- Edit distance [25]: The edit distance between an input read and, in principle, all the variants will be calculated, such that the variant with the smallest edit distance will be selected as the matched one.
- Fast matching: The tool supports also faster matching using several approximations of the edit distance. Alternatively, this matching step can be done by the user in advance. In this case the matching between reads and variants is given by fourth input file (in csv format). The set of reads which are matched to the same variant form a *variant cluster*.
3. **Alignment**: Every read is aligned according to its matched variant and an error vector is computed which represents the location and error types at each position of the variant (with insertions handled separately). Fig. 1 demonstrates an example for the alignment step.
4. **Analysis**: The matched reads and their error vectors are used in order to create error characterization and data statistics for the library, as will be described in the sequel.
5. **Report generation**: The output of our tool is a report which consists of analysis results, as selected by the user, in a customizable format.

**Figure 1:**
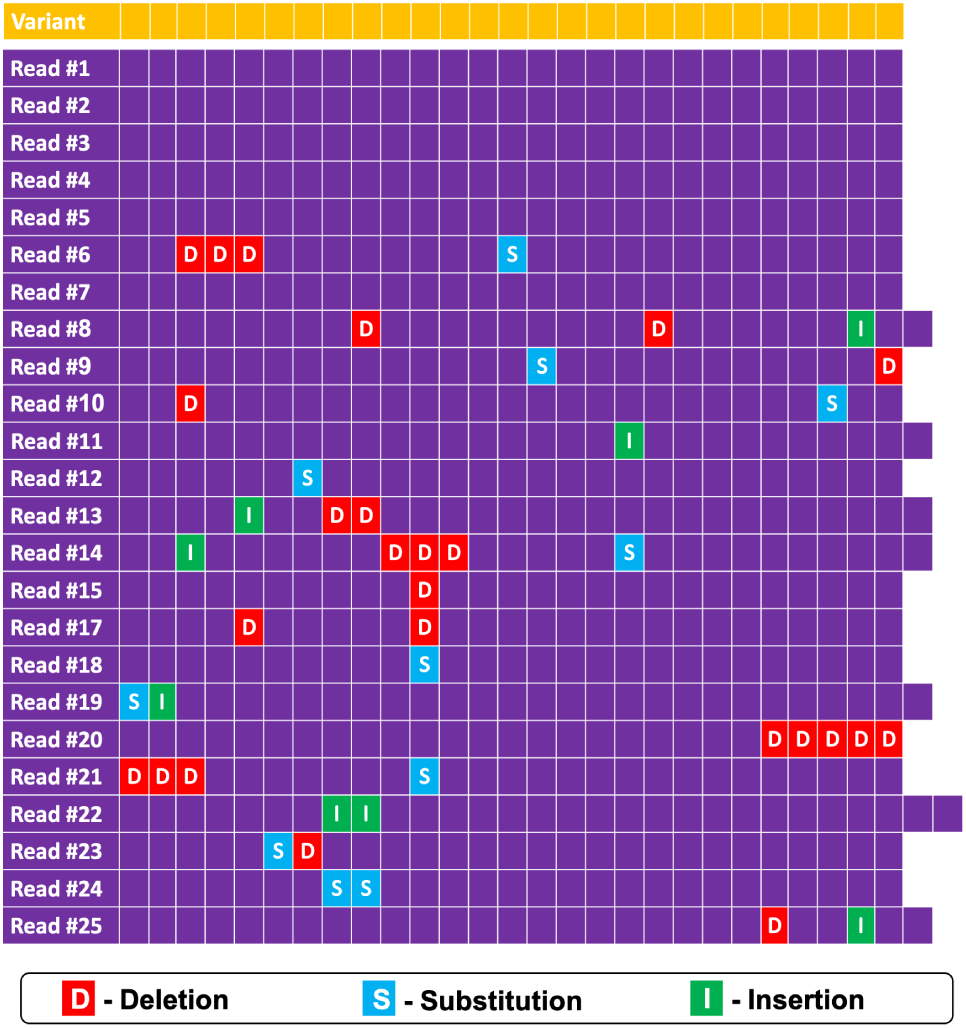
An example of 25 reads (in purple) aligned to a variant of length 27 (in yellow). For each read, the locations of the deletions, substitutions, insertions are marked in red, blue, green, respectively. This alignment output forms the basis of the analysis performed by SOLQC.

### 2.2 Statistical QC Analysis for Synthetic DNA Libraries

In this section we describe and discuss the statistical analysis performed and supported by the SOLQC tool. These statistics are explained on actual data from the experiment in [9] by Erlich and Zielinski. The details of this experiment are summarized in Table 1. These statistical results are divided into two parts; The first one addresses the composition of the synthesized library (composition statistics) and the second one addresses the errors inferred from sequencing reads (error statistics). We sampled 1,689,319 reads out of the 15,787,115 reads of the library, and analyzed only reads with length at most 4 bases shorter or longer than the design’s length, which is 152 (i.e., their base-length was between 148 and 156). Those reads were matched with their closest design variants using an approximation of the edit distance which calculated the edit distance between all reads and variants based upon the first 80 bases.

**Table 1:** Experiment by Erlich and Zielinski [9]

|  |  |
| --- | --- |
| Data size | 2.11 MB |
| Design length | 152 bases |
| Number of variants | 72000 |
| Number of reads | 15,787,115 |
| Number of sampled reads | 1,689,315 |
| Number of filtered reads | 1,427,781 |
| Synthesis Technology | Twist Bioscience |
| Sequencing Technology | Illumina miSeq V4 |

#### 2.2.1 Composition statistics

1. **Symbol statistics** (Figs. 2 and 3). This plot presents, using a stacked-bar plot, the distribution of all bases in the library by their occurrence at any position both for the reads and for the design variants. This is demonstrated in Fig. 2 for the design variants and in Fig. 3 for the reads.
  - X-axis: The position (index) in the DNA variant or read.
  - Y-axis: The number of occurrences for each base type, scaled for the sequencing depth.
  - Description: In Fig. 2, for every position (index) in the variant, the number of occurrences of each of the four bases in all of the design variants is calculated. Similarly, in Fig. 3, the number in every position is calculated according to the actual reads.
2. **Histogram of the cluster size per variant** (Figs. 4 and 5). The plot in Fig. 4 presents the histogram of the variant cluster size. That is: the number of filtered reads, per design variant.

- X-axis: The size of a variant cluster, starting from the size of the smallest variant cluster among all the variants in the library and up to the largest variant cluster value.
- Y-axis: The number of variants in the library that have a cluster of size *x*.
- Description: According to the matching step, the cluster size for each of the design variants is calculated and the histogram is generated by counting the number of variants with a given cluster size. Note that the sum of the *y* values in this histogram is the number of variants in the experiment, which is 72,000 in [9]. This plot can also have a stratified version by 5 ranges of the GC-content of the design variants, as depicted in Fig. 5. To define the 5 values of the GC-content presented in the figure, the tool takes the minimal and maximal values of the GC content as designed in the library and partitions the range between them to 5 different subranges of equal size (in terms of range). The GC-content is presented by percentage.
3. **Sorted bar plot of the number of filtered reads per variant** (Figs. 6 and 7). The plot in Fig. 6 presents a sorted bar plot for the variant cluster sizes.

- X-axis: The variant rank after sorting all variants in the library by their cluster size.
- Y-axis: The cluster size of variant *x*.
- Description: In this plot, after calculating the cluster size for each of the design variants, we sort them in a non-increasing order by the cluster size. Each variant is associated with a bar whose height corresponds to the variant cluster size. Hence, there are 72,000 bars, corresponding to the number of variants in [9]. We also plot, as shown in Fig. 7, a stratified version by 5 values of the GC-content of the variants. These 5 values of GC-content were defined by the tool as described in Fig. 5.
4. **Histogram of the length of reads** (Fig. 8). This plot presents the distribution of the different lengths of all the reads.

- X-axis: The length of the read.
- Y-axis: The number of filtered reads found in the library of length *x*, presented in log-scale.
- Description: This plot presents a histogram of the different lengths of all reads in the library.

**Figure 2:**
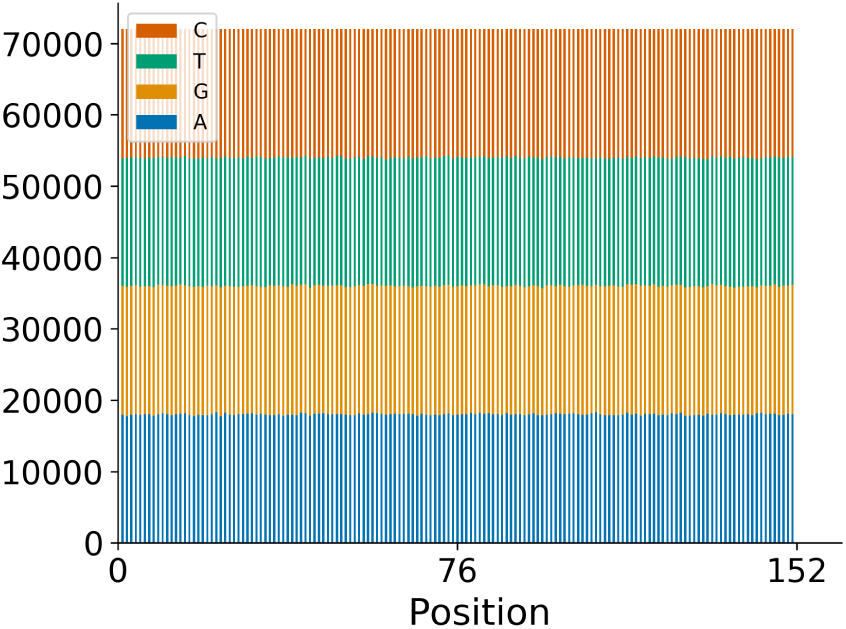
Base distribution in the design variants (see 2.2.1.1).

**Figure 3:**
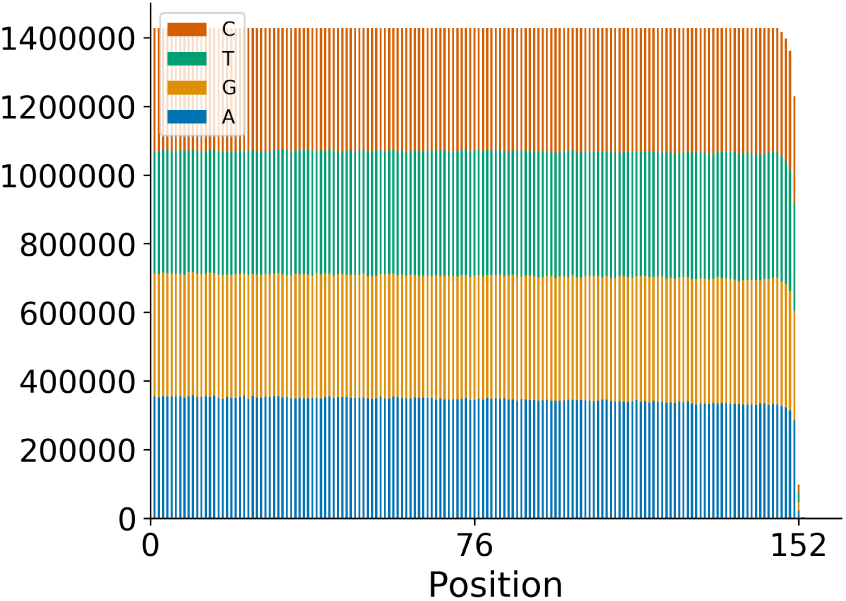
Base distribution in the reads (see 2.2.1.1).

**Figure 4:**
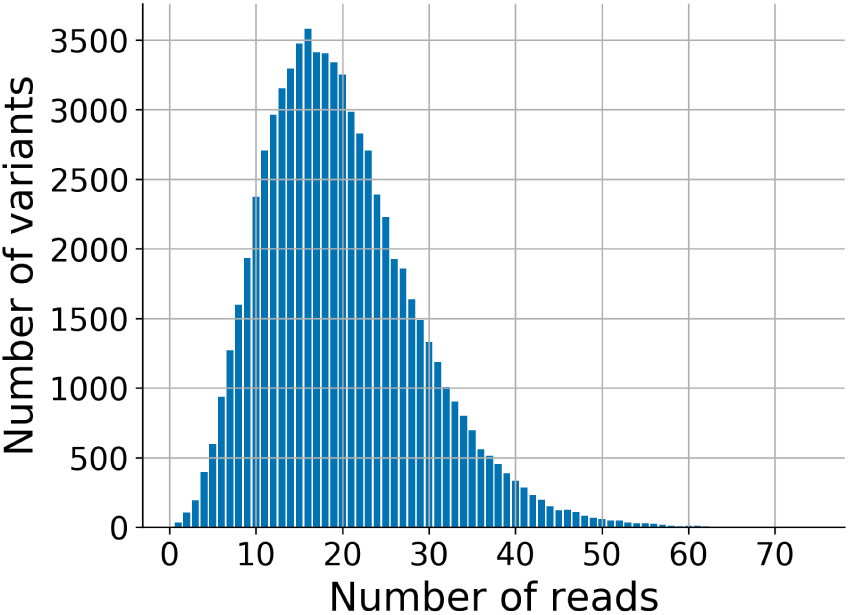
Histogram of the number of filtered reads per variant (see 2.2.1.2).

**Figure 5:**
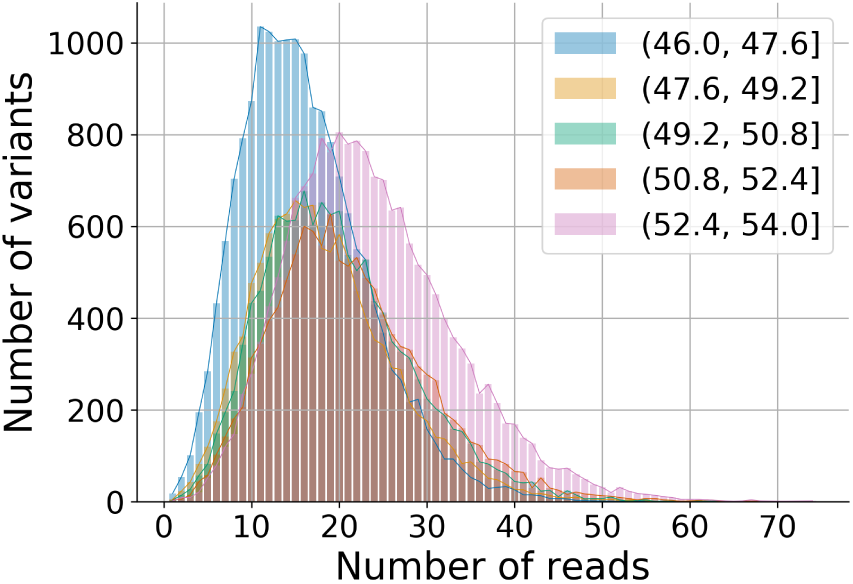
Histogram of the number of filtered reads per variant, stratified by the GC-content (see 2.2.1.2).

**Figure 6:**
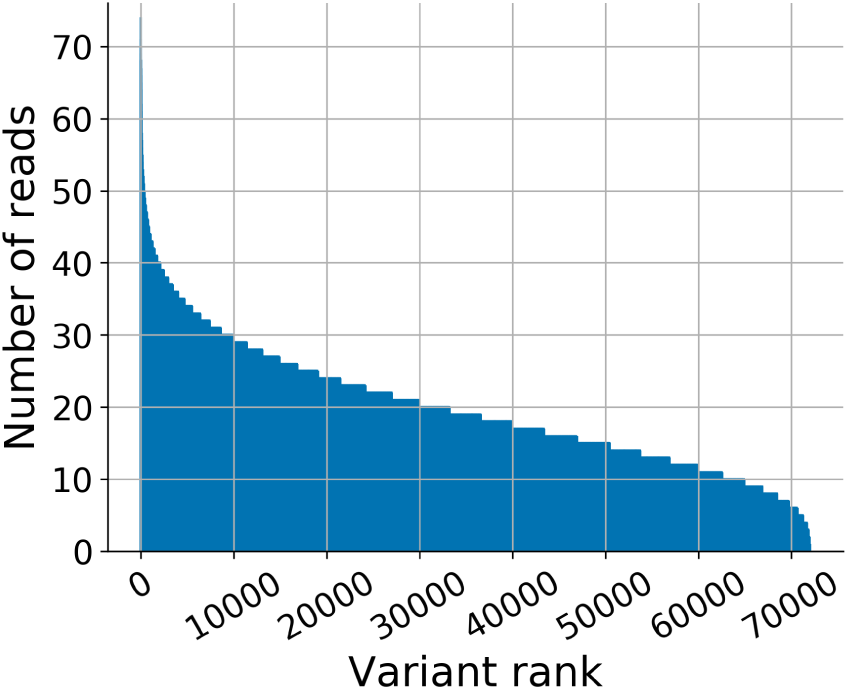
Sorted bar plot of the number of filtered reads per variant (see 2.2.1.3).

**Figure 7:**
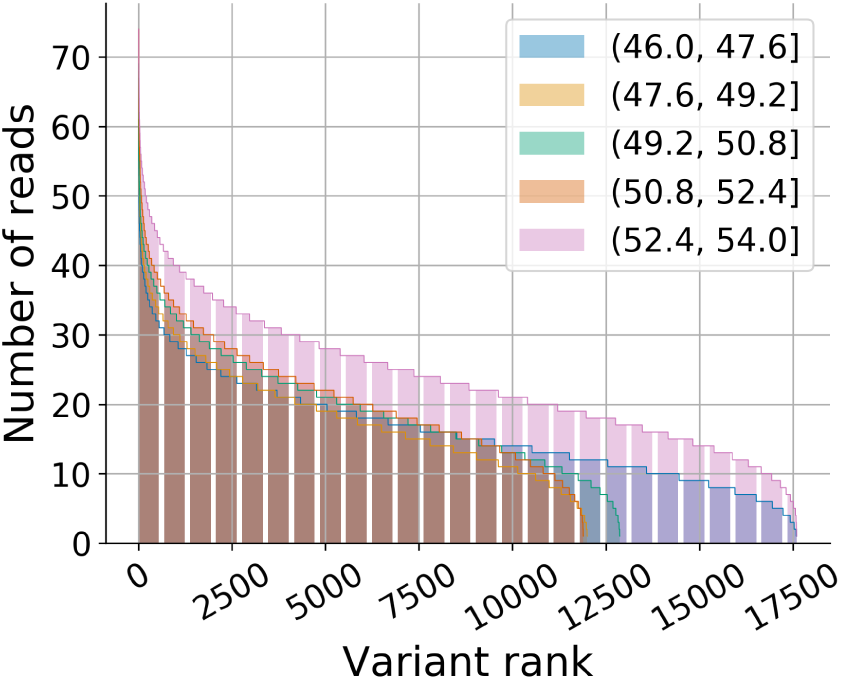
Sorted bar plot of the number of filtered reads per variant, stratified by the GC-content (see 2.2.1.3).

**Figure 8:**
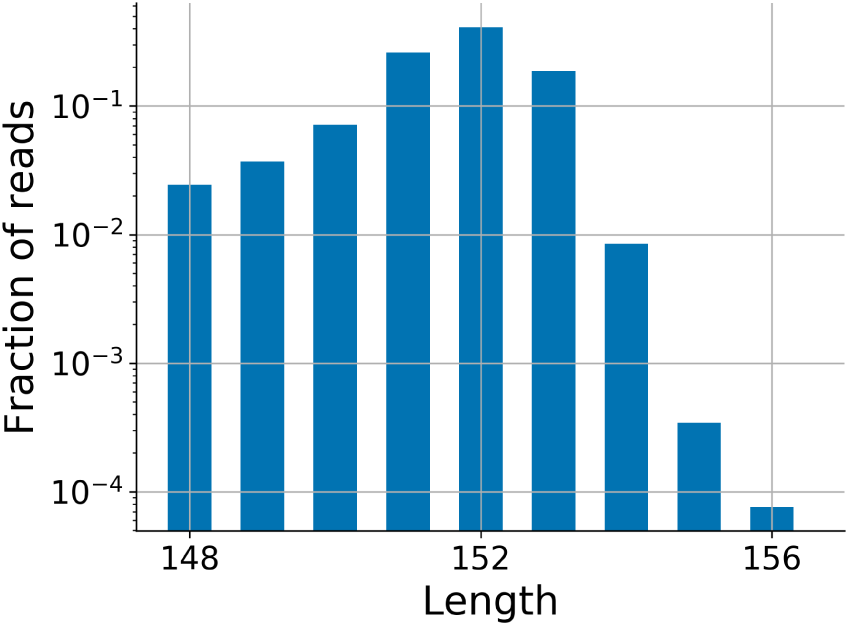
Histogram of the length of reads (see 2.2.1.4). Note that the difference between 152 and 154 is two orders of magnitude.

#### 2.2.2 Error statistics

1. **Total error rates** (Fig. 9). This plot presents the insertion, substitution, and deletion error rates as inferred from the reads in the library.
  - X-axis: Each bar presents the type of error, which can be one of the following: insertions, substitutions, single-base deletions, long deletions (deletions of more than one base), and total deletions (deletions of one or more bases).
  - Y-axis: The error rate, calculated as the ratio between the total number of errors of each type and the total number number of read bases. The plot is in log scale.
  - Description: After the alignment step, an error vector is calculated for each of the reads based upon its errors with respect to the matched variant. This error vector consists of the locations of the substitutions, insertions, and deletions in the read. See Fig. 1 for an example. For the error rates of insertions, substitutions, and deletions, we plot the ratio between the number of occurrences of each error type (in the entire sequencing data) and the total number of read bases expected in the library (number of filtered reads *×* design length^1^. For long deletions, we count each burst of at least two consecutive deletions as a single error, and then plot its ratio with the total number of read bases in the library. Lastly, the error rate of the single-base deletion is calculated is a similar way using the number of bursts of deletions of length 1.
2. **Error rate stratified by symbol** (Fig. 10). This plot presents by a heat map the symbol dependent, error distribution. Each square presents for each type of error, its error rate for the specific symbol. For insertions we address both the inserted symbol, and the symbol before the insertion. The *x*, *y* entry in the heat map is calculated to be the ratio between the number of type *y* errors of base *x* and the expected number of base *x* in the reads^2^.
3. **Error rate per position** (Fig. 11). This plot presents the error rate for every error type as it is reflected in a specific position of the strand.

- X-axis: The position in the strand, from 5’ to 3’; note that the phosphoramidite synthesis direction is 3’ to 5’. It is important to emphasize that we report rates as calculated from the alignment results. These rates reflect both synthesis as well as sequencing errors. We expect substitution and insertion errors to be primarily due to sequencing. Long deletions primarily due to synthesis.
- Y-axis: The error rates per position in all reads for single-base deletions, long deletions, substitutions, and insertions, presented in log scale.
- Description: For every position between 0 (the first position, from 5’ to 3’) and 151 (the last position in [9]) and for each error type as described in Fig. 9, the tool calculates the error rate as the ratio between the number of errors of each type and the number of filtered reads.
4. **Deletion length distribution** (Fig. 12). This plot presents the distribution of the lengths of all deletions.
  - X-axis: Deletion length, which is the number of consecutive deleted bases.
  - Y-axis: The error rate for each length of burst of deletions with exactly *x* bases divided by the number of total bases in the library.
  - Description: The tool counts the number of deletion bursts of size exactly *x* bases, based on the alignment error vector. The error rate is then calculated as the ratio between this number and the expected number of bases in the reads.
5. **Cummulative distribution based upon the number of errors** (Fig. 13). This plot presents the percentage of reads in the library with *x* or less errors.

- X-axis: Number of errors.
- Y-axis: Percentage of reads with at most *x* errors.
- Description: For a given number of errors *x*, the tool calculates the fraction of reads with at most *x* errors.
6. **GC-content error analysis** (Fig. 14). Error rates in a form of box plot based upon the GC-content. This plot depicts the reads error rates, grouped by the GC-content of their corresponding design variants. Each point represents the error rate of one of the reads in the library, with GC-content *x*. The box extends from the lower to upper quartile error rate of the reads with GC-content *x*, and plots green line at the median and green triangle at the mean value.

**Figure 9:**
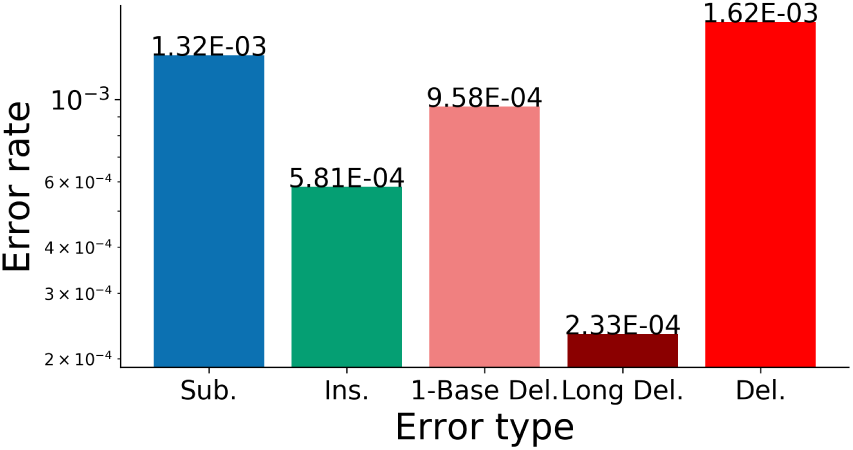
Total error rates. (see 2.2.2.1)

**Figure 10:**
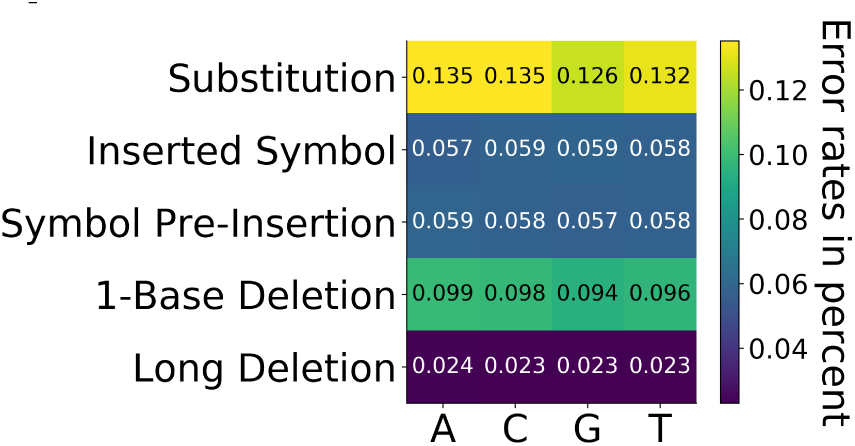
Error rates stratified by symbol. Note that the numbers are in percents. For example the value of 0.024 for “A” long deletion, means that 0.024 percents of the occurrences of base A in the library creates long deletion error. (see 2.2.2.2)

**Figure 11:**
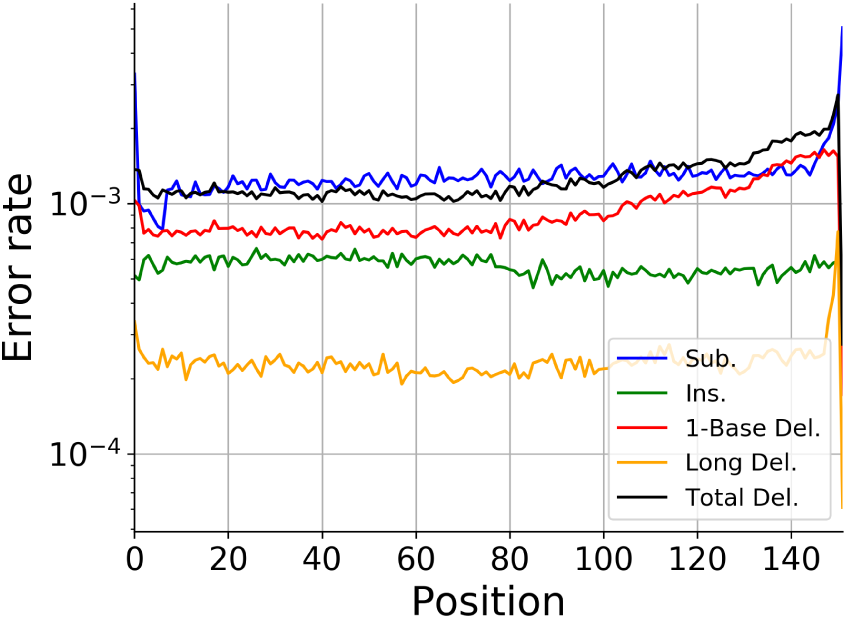
Error rates by position (see 2.2.2.3). X-axis represents position counted from the 5’ end of the designed variant. The Y-axis is logscale.

**Figure 12:**
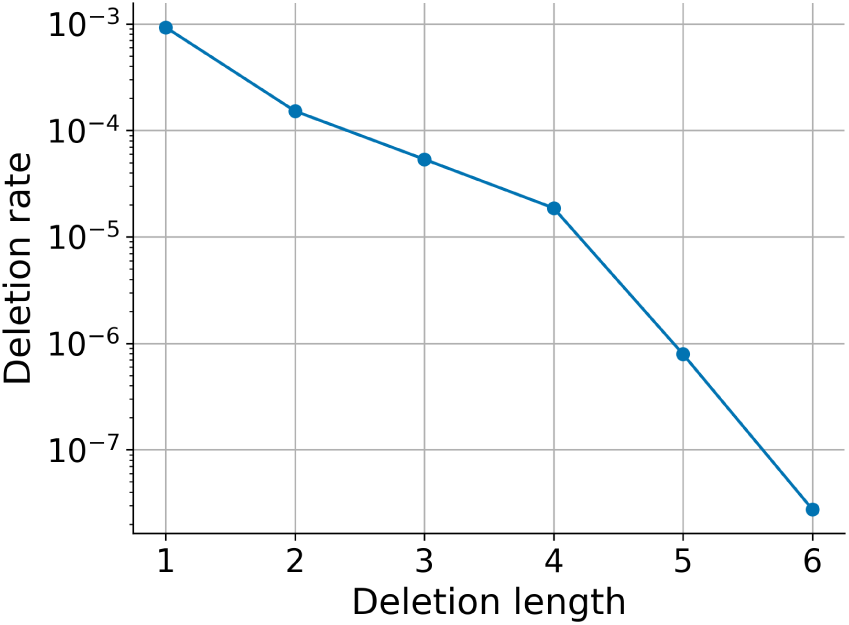
Deletion length distribution (see 2.2.2.4).

**Figure 13:**
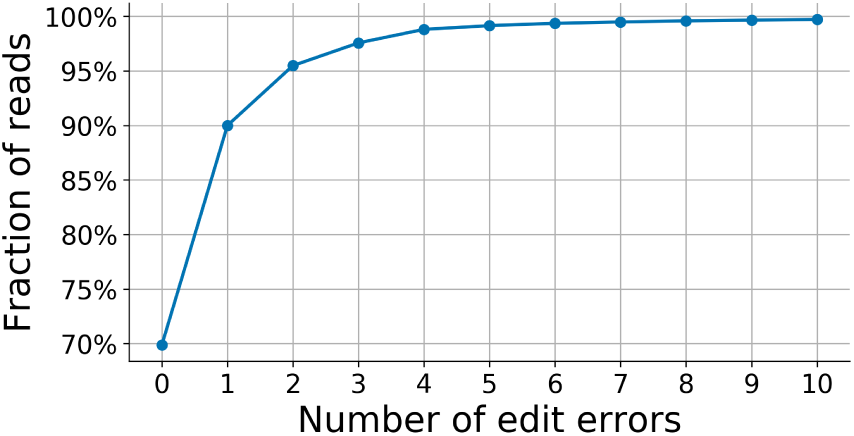
Cummulative distribution based upon the number of errors (see 2.2.2.5). Note that 70% of the reads have neither sequencing nor synthesis error.

**Figure 14:**
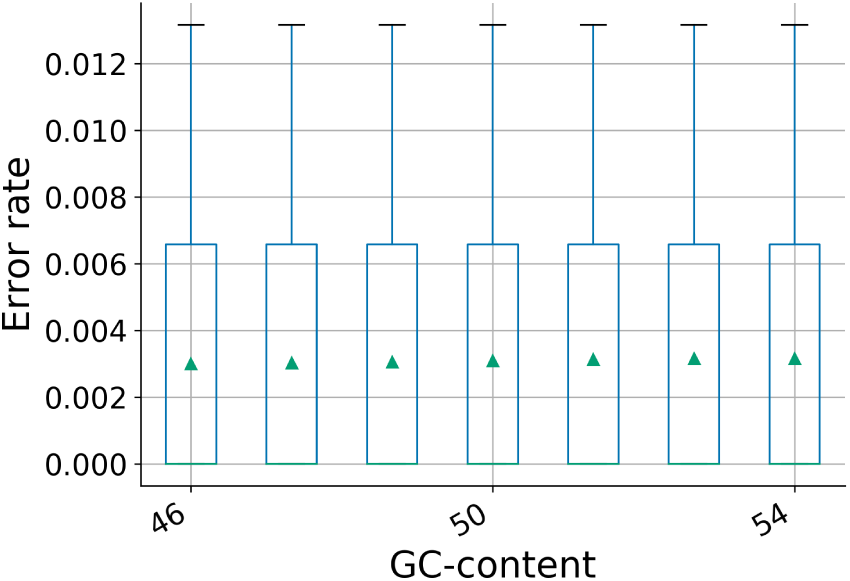
Error rates by the GC-content (see 2.2.2.6).

## 3 Results

In this section, we present several results from the analysis of four synthetic DNA libraries. These results are based on previous experiments for storage applications conducted by Erlich and Zielinski [9], Grass et al. [12], Organnick et al. [20], and Yazdi et al. [27]. While OLs are used for a variety of applications, we focused on data storage OLs as the library data for these is typically more accessible. We matched each read with its relevant variant using edit distance estimation as will be described below. The analyzed data sets and their details are summarized in Table 2. We next present how we process the data of each experiment.

**Table 2:** Syntheti DNA Libraries

|  | EZ-17 [9] | G-15 [12] | O-17 [20] | Y-16 [27] |
| --- | --- | --- | --- | --- |
| Data storage size | 2.11 MB | 83 KB | 200 MB (9.503066 MB) | 3,633 bytes |
| Design length (bases) | 152 | 158 | 150 | 880-1,060 |
| Number of variants | 72,000 | 4,991 | 607,150 | 17 |
| Number of reads | 15,787,115 | 3,312,235 | 62,879,612 | 6,660 |
| Number of filtered reads | 1,427,781 | 1,945,744 | 91,898 | 6,660 |
| Synthesis Technology | Twist Bioscience | CustomArray | Twist Bioscience | Integrated DNA Technology (IDT) |
| Sequencing Technology | Illumina miSeq | Illumina miSeq | Illumina NextSeq | MinION |

### 3.1 Pre-processing and Filtering of the Libraries

- Erlich and Zielinski [9], will be referred in this paper as EZ-17: As explained in Section 2.2, we analyzed this library using a sample of 1,689,315 reads out of the reported 15,787,115 reads. In this library the design length of each variant was 152. We present example results from three different filtering schemes:

1. Filtering only reads with length between 148 and 156 - 1,427,781 reads.
2. Filtering only reads with length between 142 and 162 - 1,466,069 reads.
3. Analyzing all the reads in the sample - 1,689,315 reads. The estimated matching between each of the reads and its design variant was calculated in two steps. First, the edit distance of the first 80 bases between the read and each of the variants was calculated. Then, the read is matched with the closest variant according to this calculation.
- Grass et al. [12], will be referred in this paper as G-15: The analysis of this library is based on all of the 3,312,235 reads. The length of each variant in this library was 158, with two primers of length 20 at the 5’ end and 21 at the 3’ end. The results presented were calculated according to the 117 bases of the data in each of the reads. The reads were filtered by their length: 1,945,744 reads with length between 112 to 122 bases were analyzed by the tool. The estimated matching of the reads to their corresponding design variants in G-15 was performed as in EZ-17.
- Organick et al. [20], will be referred in this paper as O-17: The analysis of this library is based on a sample of 101,243 out of the 62,879,612 reads of one file of the library. The design length of each variant in this library was 150. Similarly to G-15 [12], there were two primers of length 20 at each end. Hence, the reads were filtered by their length: we omitted the primers from each read, and analyzed 91,898 reads with length between 105 and 115 bases. The results are presented for the information bases (the primers were trimmed). The estimated matching of the reads in O-17 was performed as in EZ-17.
- Yazdi et al. [27], will be referred in this paper as Y-16: The results presented are based on all the 6,660 reads in the library. This library consists of 17 variants - 15 of length 1,000, one of length 1060 and one of length 880. The estimated matching of the reads to their corresponding design variants, was done in a similar way that used for EZ-17. However, since the number of variants was significantly smaller, we were able to calculate the edit distance for the entire strand between each read and all of the variants. Then, the read was matched with the closest variant. In this experiment, the design variants were longer and the reads were sequenced by the MinION sequencing technology. Hence, this data is likely have different error characteristics than those observed for the other three.

### 3.2 Analysis of Synthetic DNA Datasets

1. **Total error rates**. The results show significant differences between the four experiments. The three experiments of EZ-17 [9], G-15 [12], and O-17 [20] show higher rates for deletions and substitutions than insertions. EZ-17 and O-17 have the lowest error rates overall. In Y-16 [27], we observe higher rates for insertions rather than deletions and substitutions. Moreover, the error rates in Y-16 [27] are higher by two orders of magnitude than the other three. These results are presented in Fig. 15.
2. **Cummulative distributions based upon the number of errors**. As mentioned above the data of Y-16 [27] is much more erroneous. Indeed, we can see that none of its reads had less than 100 errors. In EZ-17 [9] and O-17 [20], 70% and 60% of the reads were synthesized and sequenced without any error respectively, while only 30% of the reads in G-15 [12] show no errors at all. These results are presented in Fig. 16.
3. **Error rates, stratified by symbol**. Base G showed slightly less errors compared to the other bases in EZ-17 [9]. Similarly, base A and base G showed slightly lower error rates compared to the other bases in Y-16 [27]. However, in G-15 [12], and in O-17 [20], base C was the least erroneous base. These results are presented in Fig. 17.
4. **Histograms of the length of the reads, using different filtering schemes in EZ-17 [9]**. We can see that in each of the filtering schemes we used, the length of the majority of the reads was 152 (the designed length) or shorter. These results correspond to our findings that deletions were the most dominant errors in the library. These results are presented in Fig. 18.
5. **Histograms of the cluster size per variant**. Fig. 19 shows the distributions of the cluster size per variant for the experiments in EZ-17 [9] and G-15 [12]. While the shape of the distribution of EZ-17 [9] has the form of a normal distribution, there is no similar trend in G-15 [12]. However, it is possible to notice that the number of variants decreases with the size of the cluster size. Furthermore, in G-15 [12] we also observed very large clusters of size ranging between 2,000 and 8,000, which were omitted from the figure for its clarity.
6. **Error rates by GC-content**. The results (presented in Fig. 20) show that the median and the mean values of the read error rates increase with its designed GC-content in G-15 [12]. In Y-16 [27] we can surprisingly see error rates which are greater than 1. Such high error rates are encountered when there is a large number of insertions together with deletions and substitutions in the read such that the number of errors is strictly larger than the design length. These results corresponding to our findings that insertions were the most dominant errors in this library.
7. **Error rate per position**. In all four experiments analyzed, the error rates in the 3’ end are greater than the error rates in the 5’ end. Note that in Y-16 [27] there were different design lengths: 880, 1,000 and 1,060. For uniformity, we present only results of reads which correspond to variants of length 1,000. These results are presented in Fig. 21.

**Figure 15:**
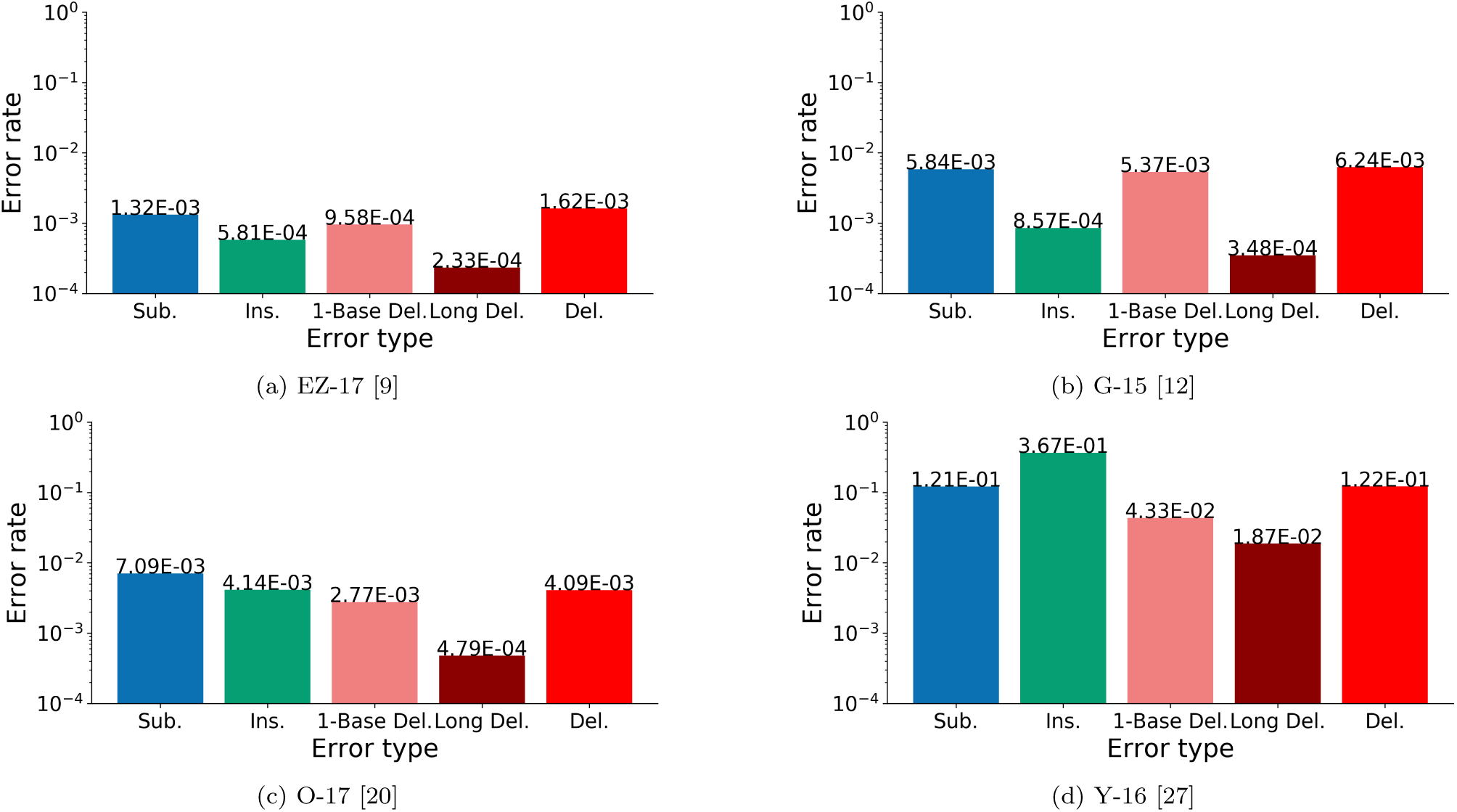
Total error rates in the four datasets. Note that total error rates are also due to sequencing errors. While EZ-17 [9], G-15 [12], and O-17 [20] were sequenced on Ilumina sequencing machines, the fourth dataset, Y-16 [27], used a much noisier sequencing platform, which explains the differences in the error rates.

## 4 Use-Case Examples

In this section we present several use-case examples for SOLQC.

a. **Design-quality evaluation**. Different libraries can have different robustness levels. For example: a design of one library can have many homopolymers, while another can limit the presence of homopolymers. SOLQC outputs statistical reports describing the error behavior of a given library/design. The user can create several small test experiments with different designs and properties. Then, the user can use SOLQC to evaluate the effect of different designs on the error behavior. This analysis can then be considered as part of the final design of the library.
b. **Binning of synthetic DNA-libraries**. The result of a sequencing reaction on a given library does not include the matching of each read to its design variant. SOLQC provides several methods to bin the reads according to their corresponding design variants. The matching/clustering methods can be performed on libraries with or without the barcode. In addition, users can get coverage depth statistics from SOLQC as well as quality related statistics, which can be different for different variants or set of variants. Lastly, in applications like data storage, the set of reads that is binned to any given variant can be used in order to decode the stored variant.
c. **Comparison of different synthesis and sequencing technologies**. SOLQC provides its users composition and errors statistics. Accordingly, the user can synthesize libraries using several synthesis technologies and their process parameters. Then, the user can compare the quality of the results of each technology and/or of each parameter configuration. In order to optimize the process parameters, the experiments can be conducted with the same design while using different parameter configuration. Thus, it is possible to determine how to choose the best configuration.
d. **Design of error-correcting codes and coding techniques for DNA-storage**. In data storage applications, SOLQC can be used as a characterization tool of the DNA channel. The user can characterize the DNA channel using data from previous experiments of various technologies and design parameters. Then, using this information, the user can design appropriate error-correcting codes and coding techniques to improve the error rates.
e. **Standardization and reproducibility**. SOLQC enables determining whether a library is behaving as previous libraries from the same vendor with similar preparation characteristics. This enables comparison between the same library preparation protocol performed in different labs, or in the same lab at different times or by different lab members.

**Figure 16:**
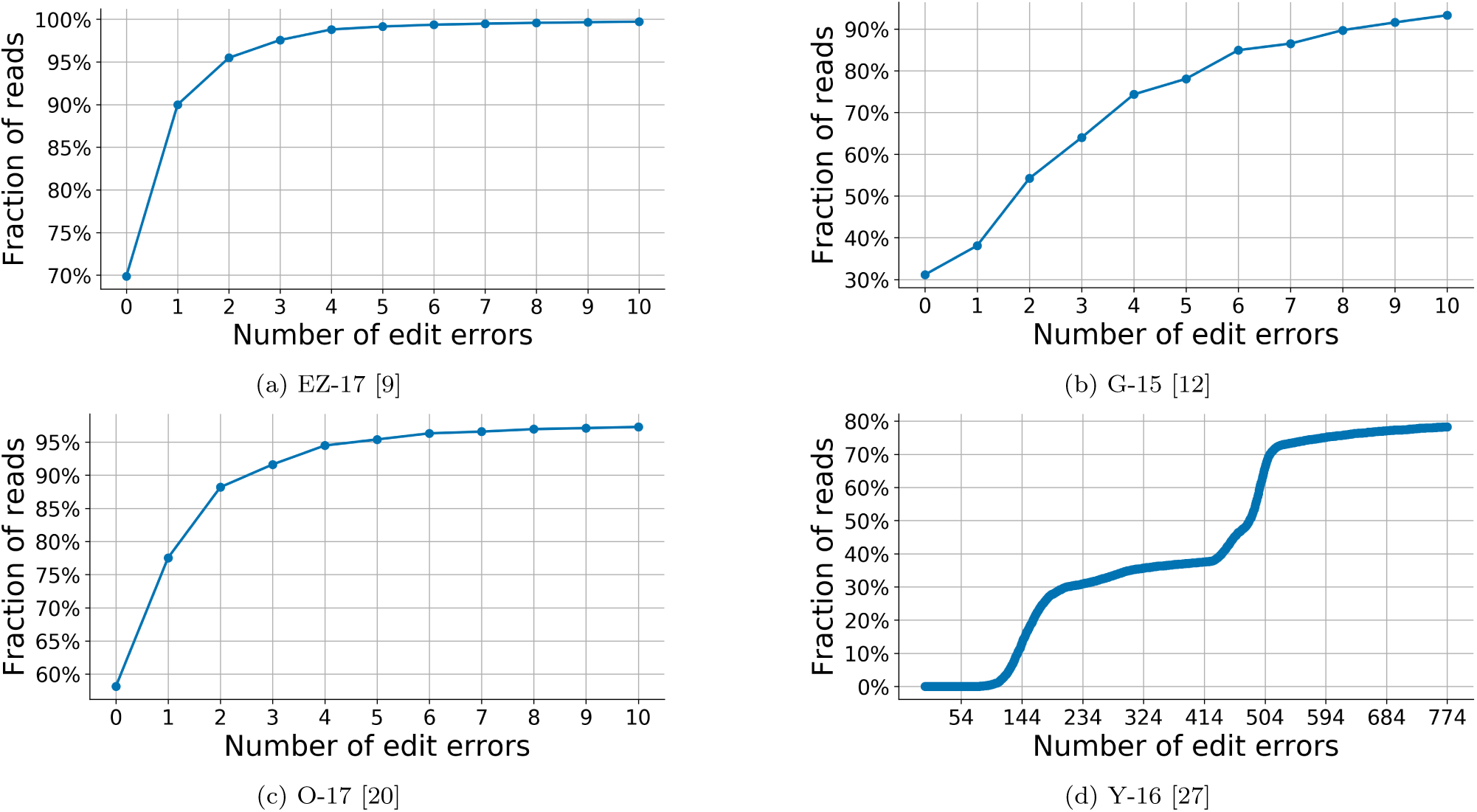
Cummulative distributions based upon the number of errors. Note differences in the range of Y-axes in the four figure. Also note that the X-axes are truncated (see 2.2.2.1).

## 5 Discussion

In this work we presented SOLQC, a software tool designed to characterize synthetic DNA libraries. While SOLQC provides many useful tools and features, it will benefit from further development in several aspects:

a. **Matching approximations**. Matching each read with its design variant is a complex calculation, especially when the library is not barcoded and/or when there are many variants in the library. In fact, the matching step is the heaviest step in any initial run of the SOLQC pipeline. Hence, we plan to provide, in the future, several faster approximation approaches to the matching step.
b. **Reconstruction algorithms**. When synthetic DNA libraries are used for data storage applications, the first step of decoding the data is to reconstruct the original variant out of the noisy reads. We plan to add to SOLQC additional features related to this step. Thus, SOLQC will perform reconstruction on a given cluster of reads, in order to decode the sequence of the original variant.
c. **Additional statistics**. SOLQC will report several more statistics in the future. These statistical analysis will examine more deeply whether there is a connection between the characteristics of the design variants and the errors observed for them.

**Figure 17:**
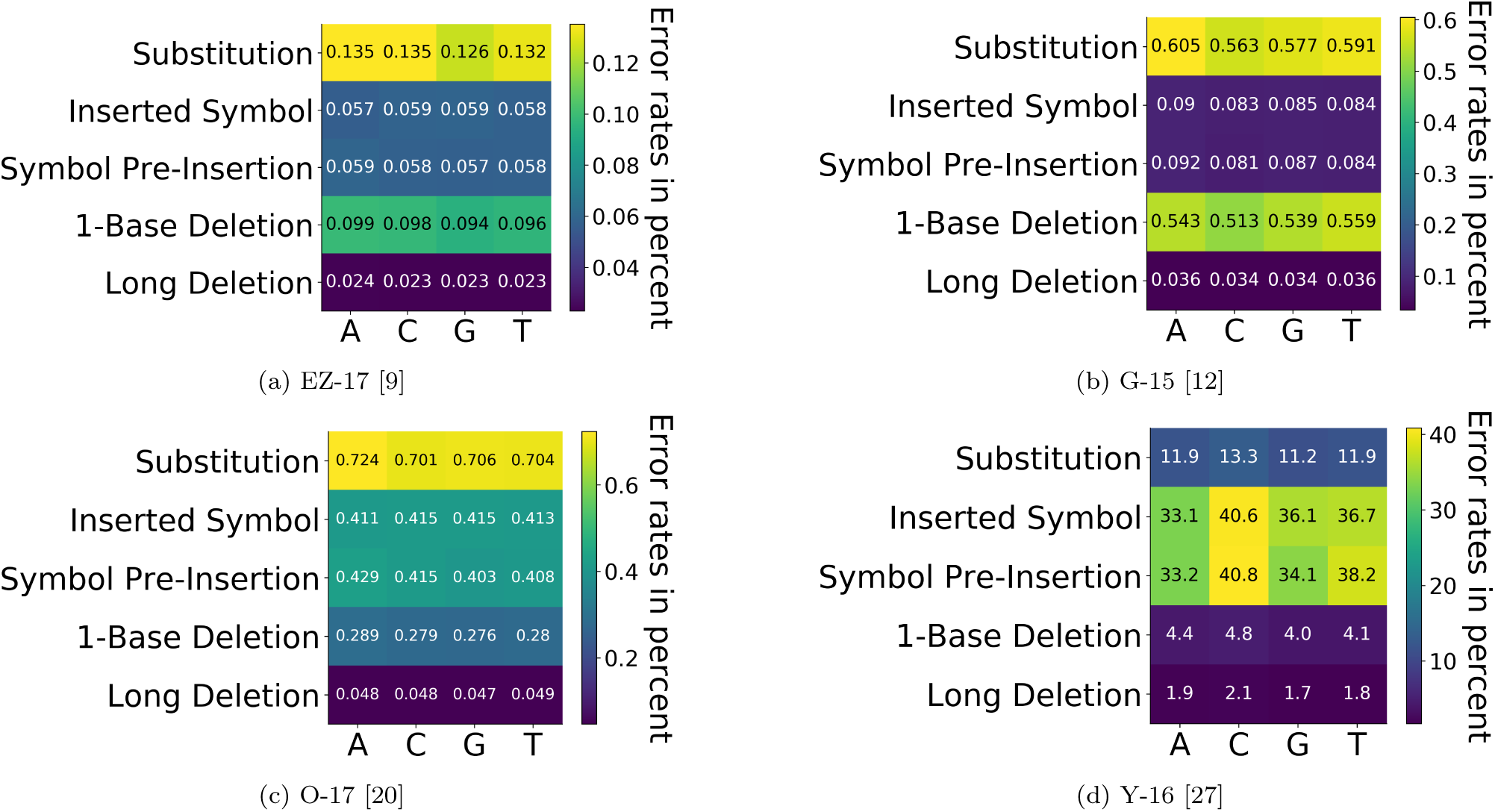
Error rates, stratified by symbol in the four datasets (see 2.2.2.2). Note that the colorbars are different in each plot.

**Figure 18:**
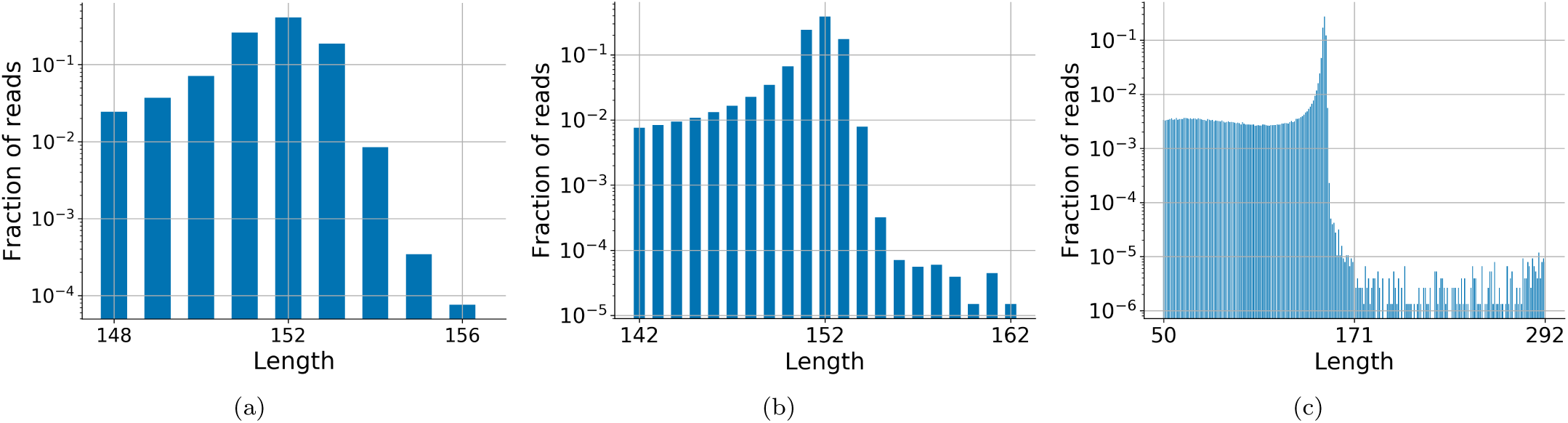
Histograms of the length of the reads in EZ-17 [9], using different filtering schemes (1), (2), and (3), respectively (see 2.2.1.4).

**Figure 19:**
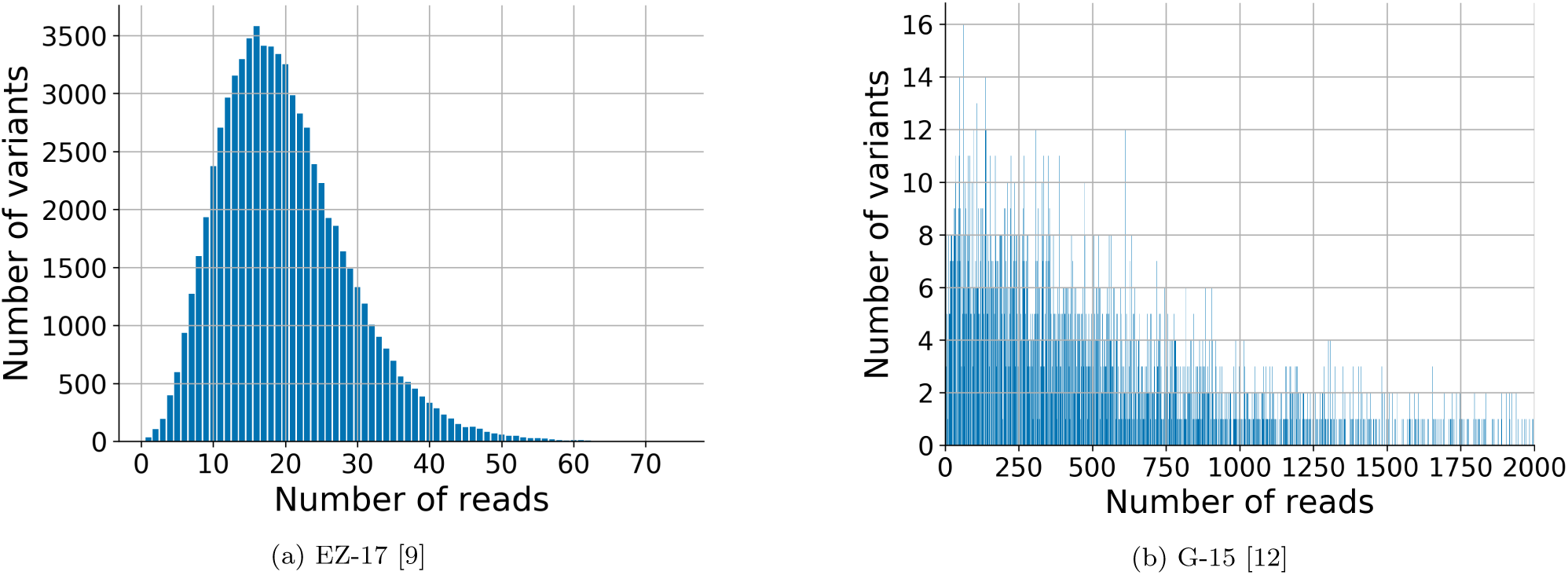
Histograms of the cluster size per variant in two of the four datasets (see 2.2.1.2).

**Figure 20:**
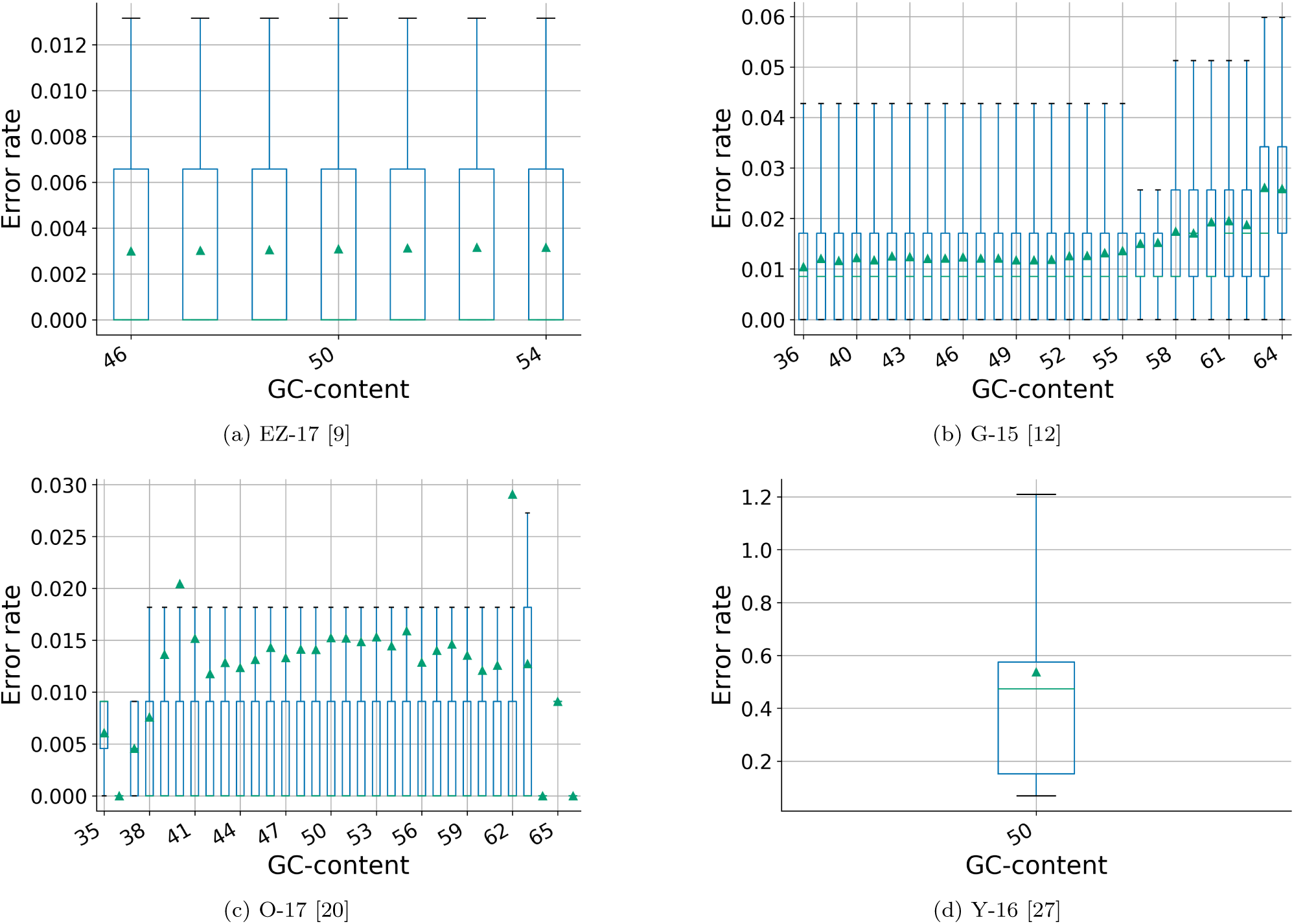
Error rates by the GC-content in the four datasets. Note the differences in the range of the Y-axis in the four figures (see 2.2.2.6).

**Figure 21:**
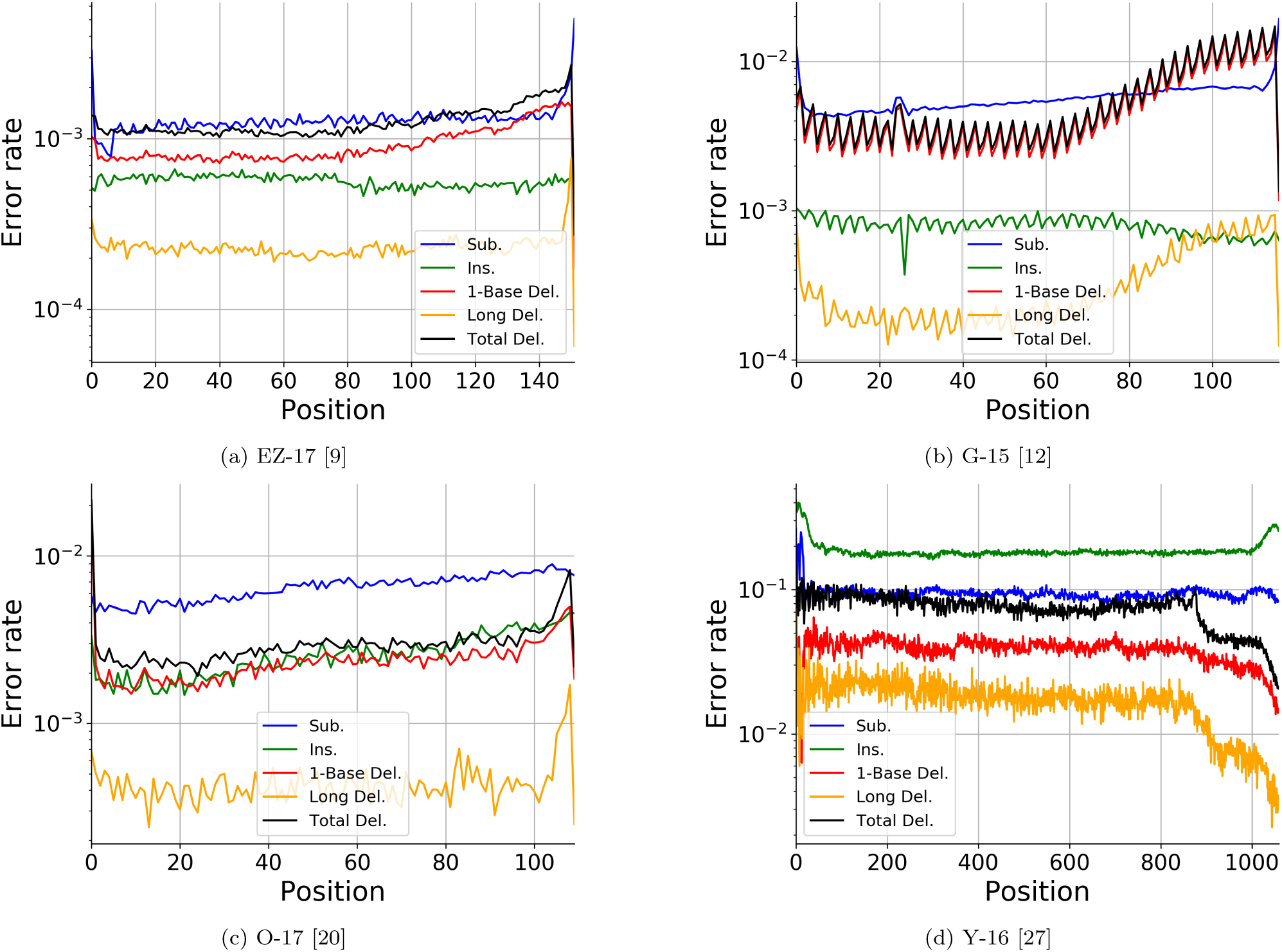
Error rates per position in the four datasets. X-axis represents position counted from the 5’ end of the designed variant, note the differences in the range of the X-axis and Y-axis in the four figures (see 2.2.2.3).

## Supporting information

An example of a report generated by SOLQC.

## 6 Installation Link

https://yoav-orlev.gitbook.io/solqc/.

## 7 Acknowledgement

We thank the authors of [9, 12, 20, 27] for sharing and providing the data of their DNA-storage experiments to this paper. We also thank Hossein Yazdi, Lee Organick, Karin Strauss, and Yaniv Erlich for helpful discussions. We thank the Yakhini research group for useful discussion and comments. We thank Roee Amit and his lab, especially Sarah Goldberg, for meaningful discussion and useful feedback. Finally, we thank Matilda Lidgi, Danit Goldberg, Amir Biran, Alex Yucovich and Batel Carmona for their great contribution to this work. This project has received funding from the European Union’s Horizon 2020 Research And Innovation Programme under grant agreement No. 851018.

## Footnotes

1 For example: the deletion rate in Fig. 1 is 24/(25×27), which is calculated to be the ratio between the number of red squares (24) and the product of the number of rows (25) with the variant length (27).

2 The expected number of base *x* in the reads is calculated as the sum of the products of the number of base *x* in each of the design variants, and the number of reads matched to it.

