## Supplementary material for "SOLQC : Synthetic Oligo Library Quality Control Tool": An example of a report generated by SOLQC.

### Analysis Pipeline

Started with 1427781 reads

Number of reads after pre-processing = 1427781

Fraction of sequences who passed pre-processing = 1.00

Number of matched reads = 1427781

Fraction of sequences who got matched from pre-processed= 1.00

Fraction of sequences who got matched from all reads= 1.00

Symbol statistics by design

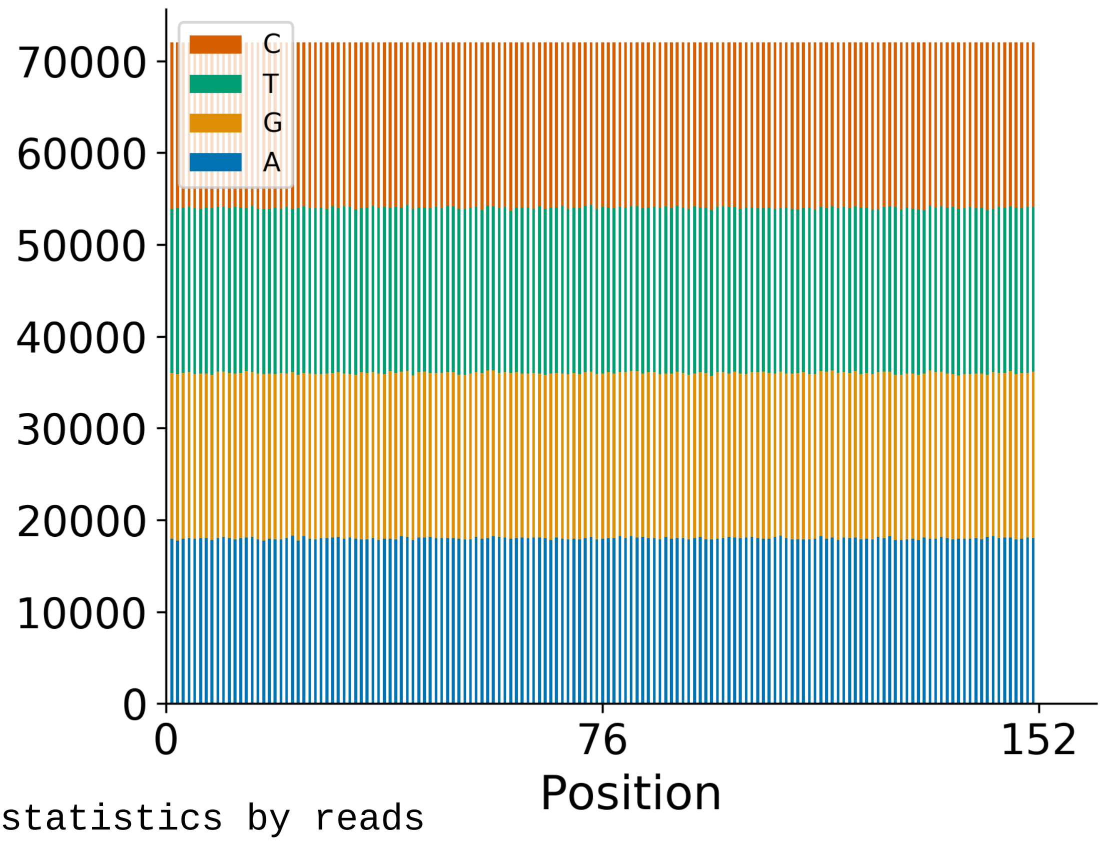


Symbol statistics by reads

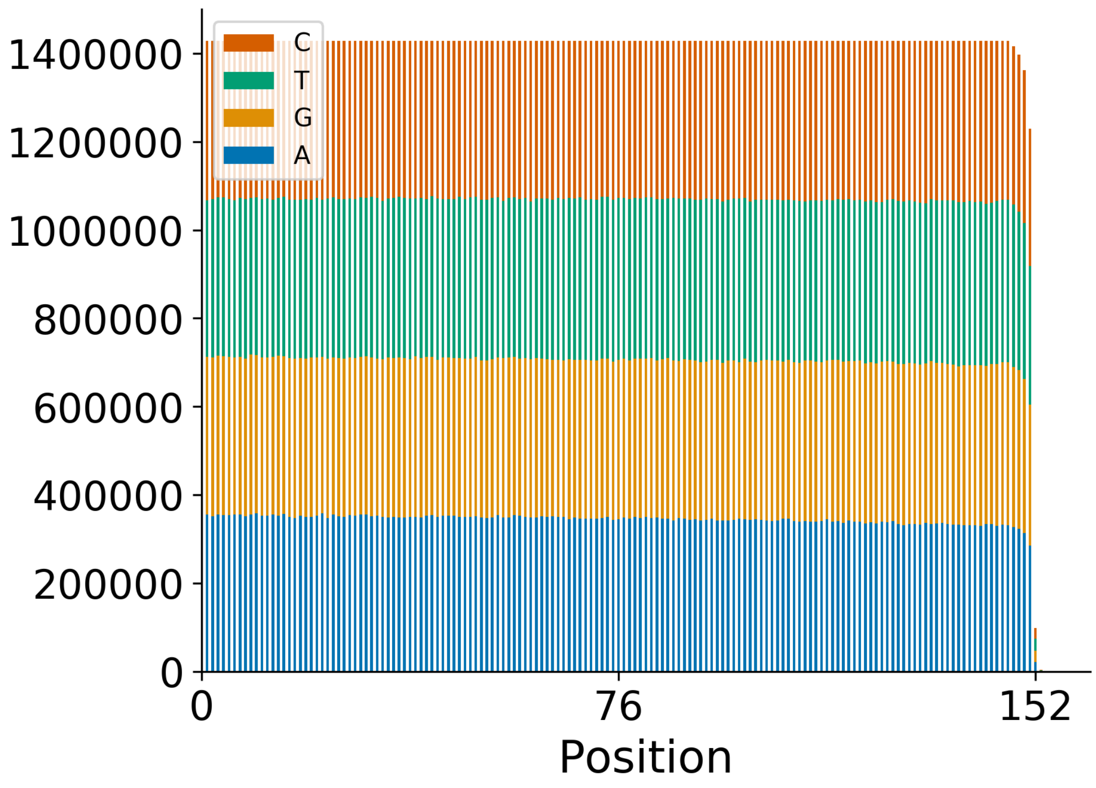

Sorted bar plot of the number of filtered reads per variant

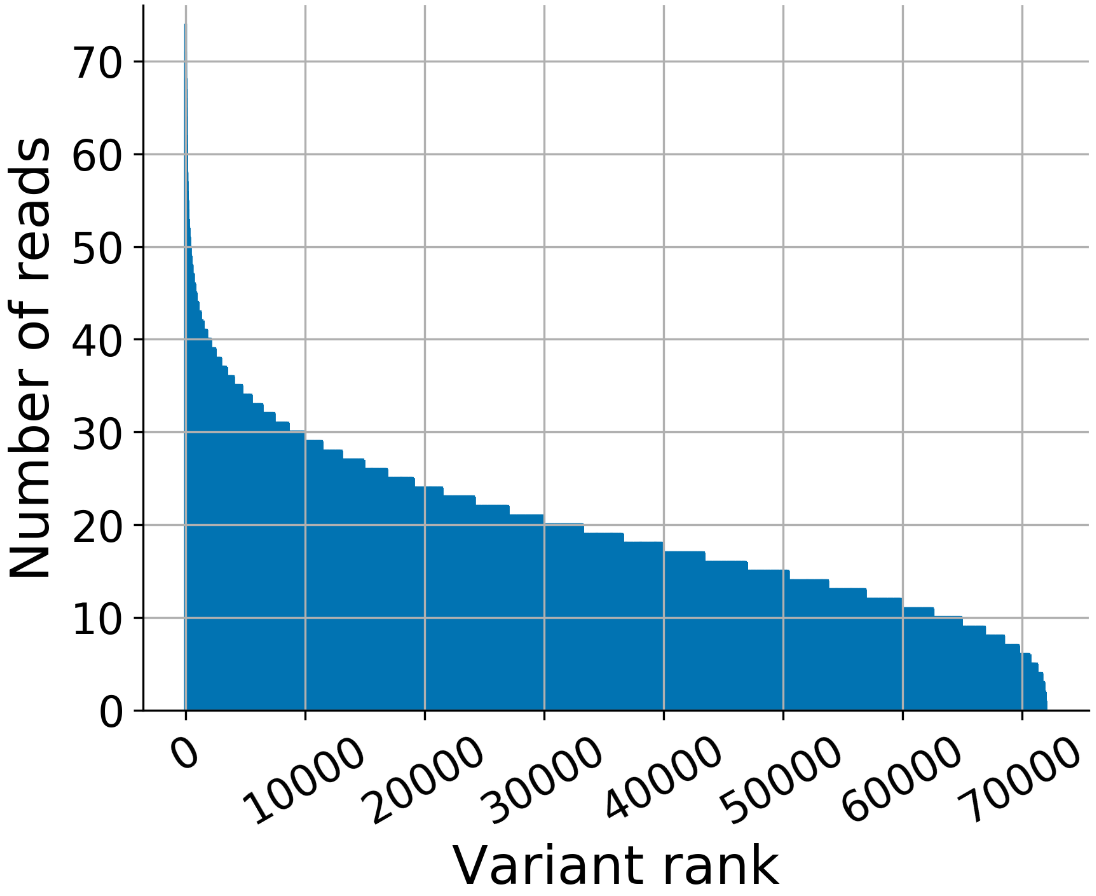

Histogram of the number of filtered reads per variant

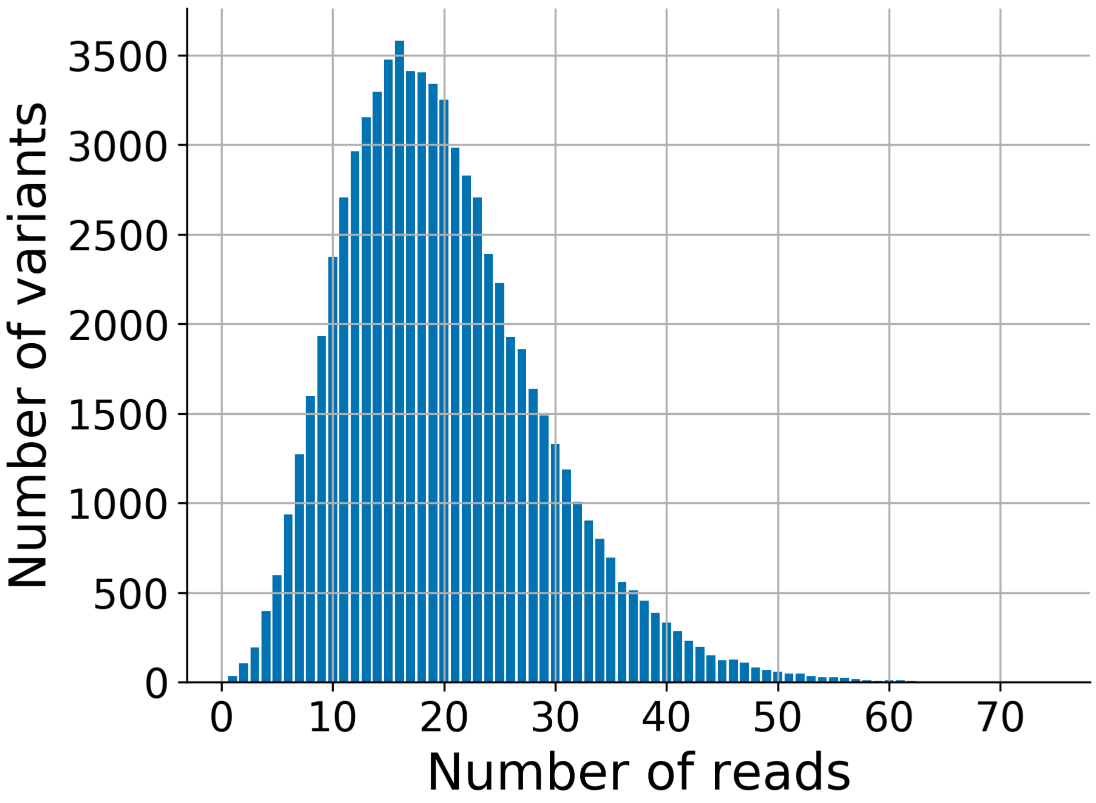

Sorted bar plot of the number of reads per variant, stratified by GC

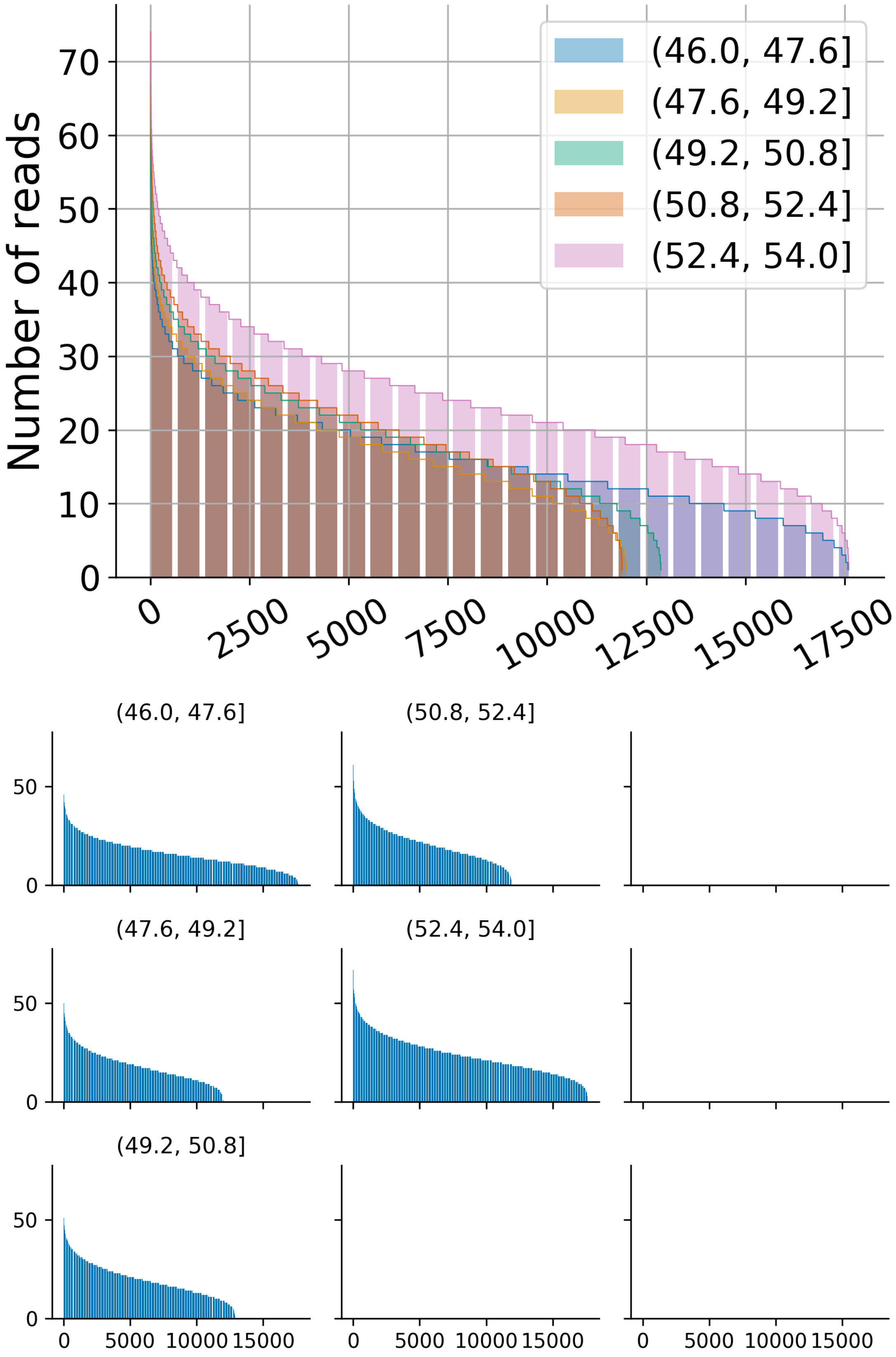

Histogram of the number of reads, stratified by GC Content

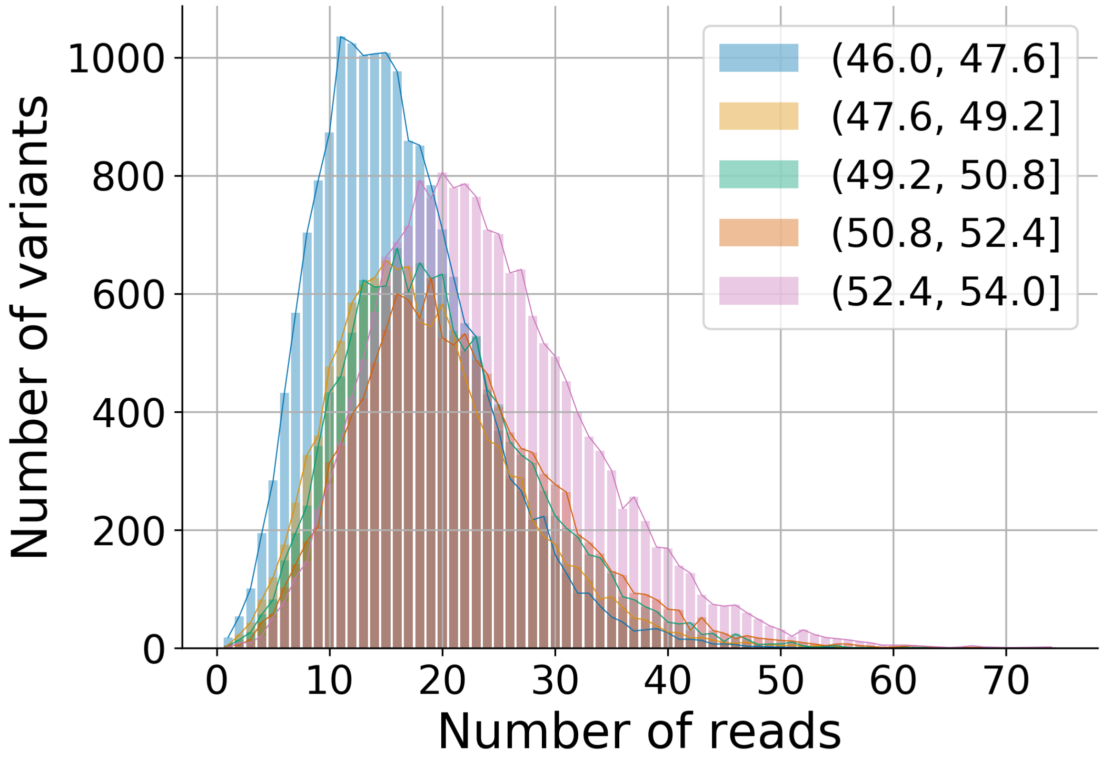


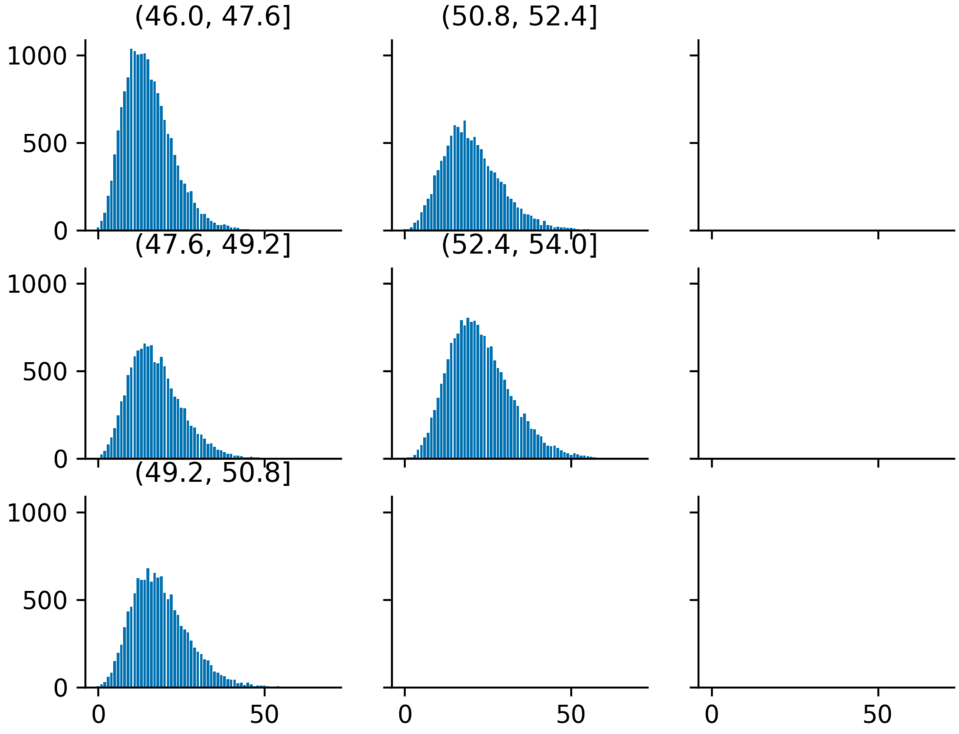

Reads length distribution

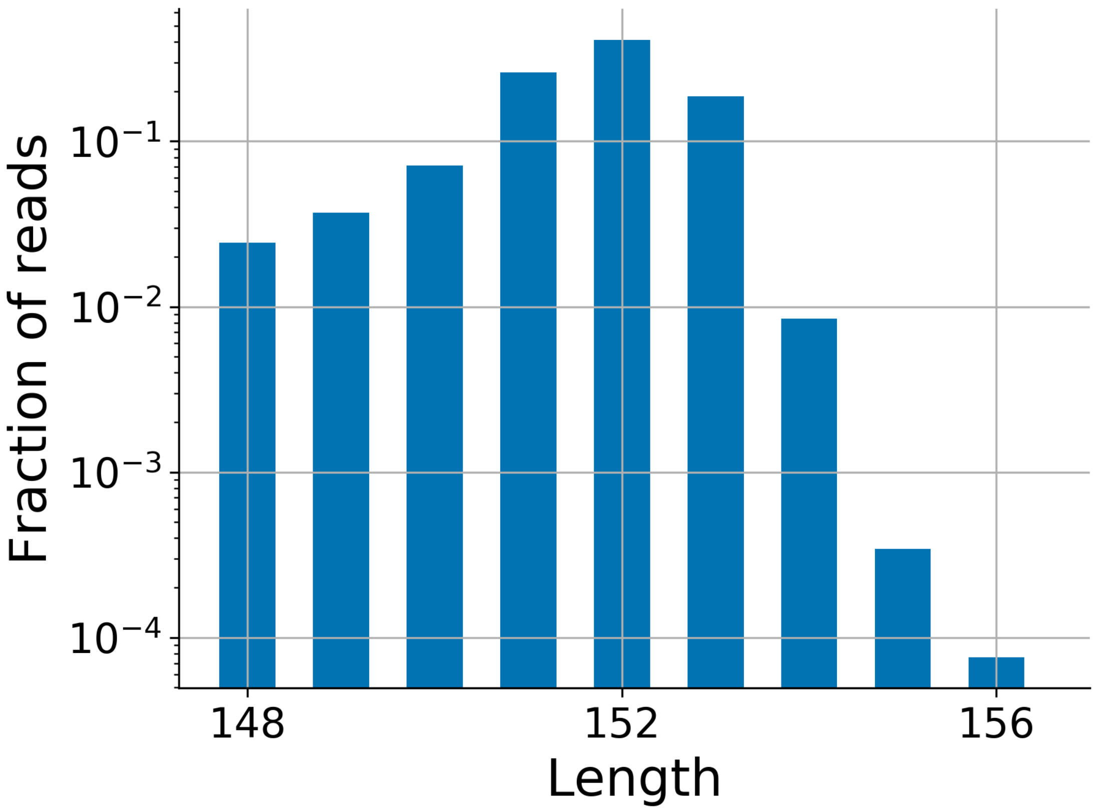

Error rates analyzer

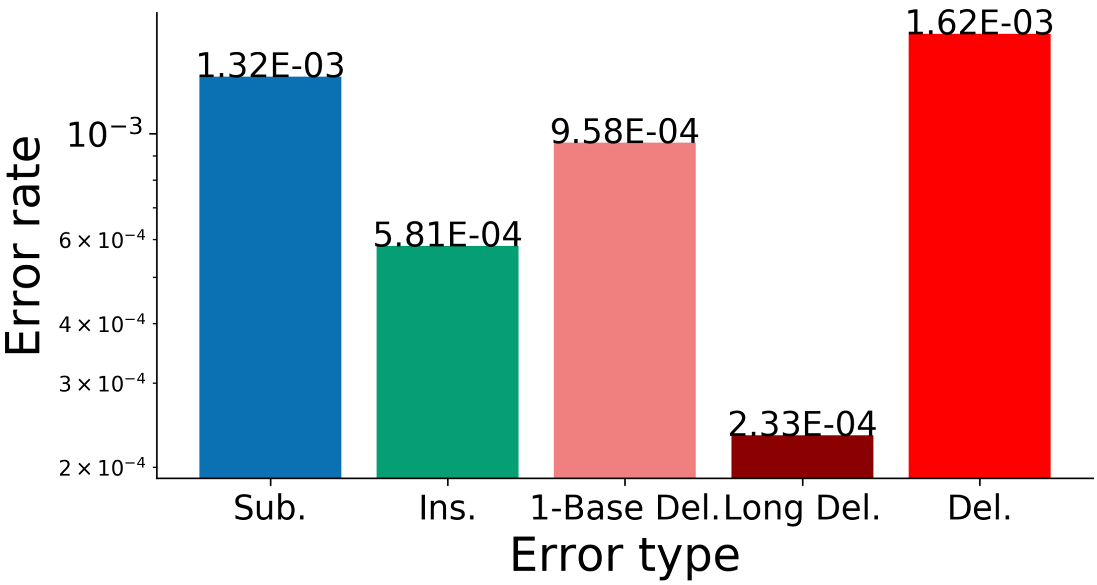

Error rates, stratified by symbol

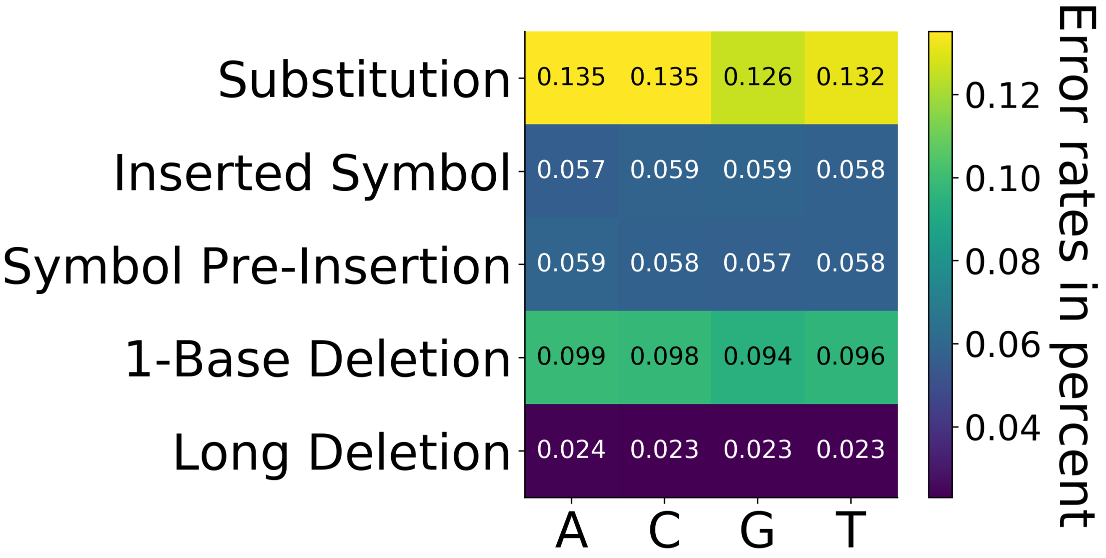

Error rates per position (5' to 3')

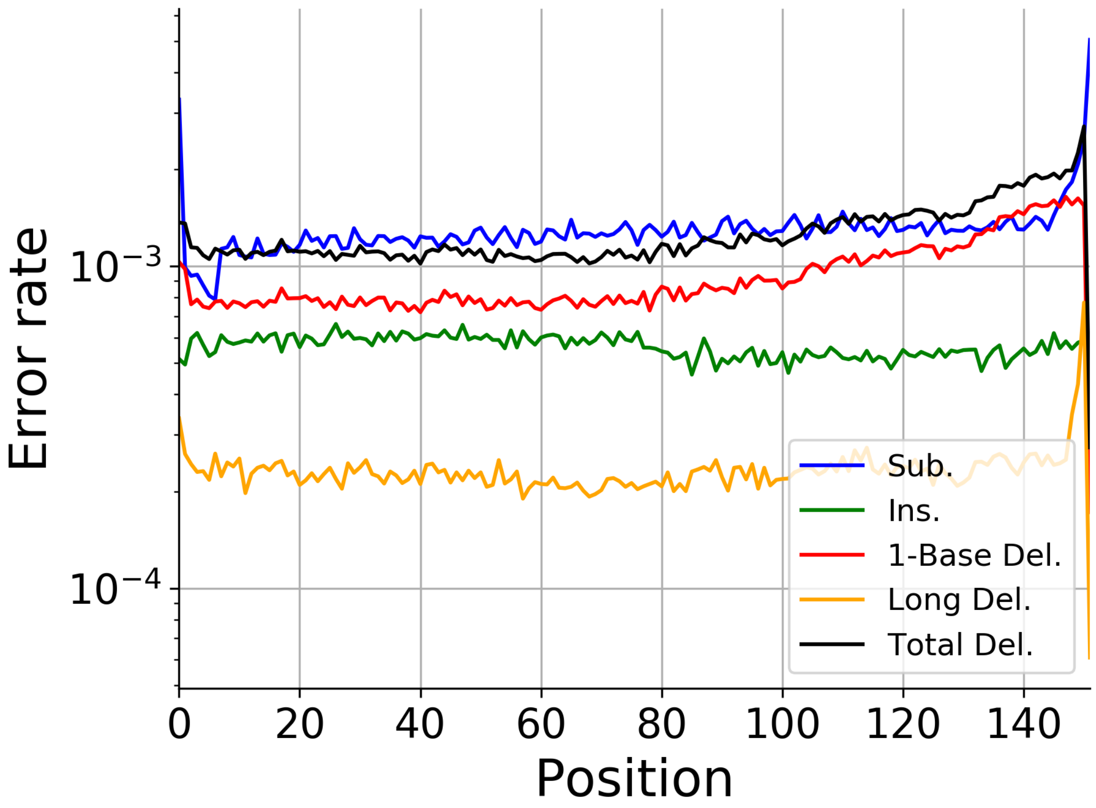

Deletion length distribution

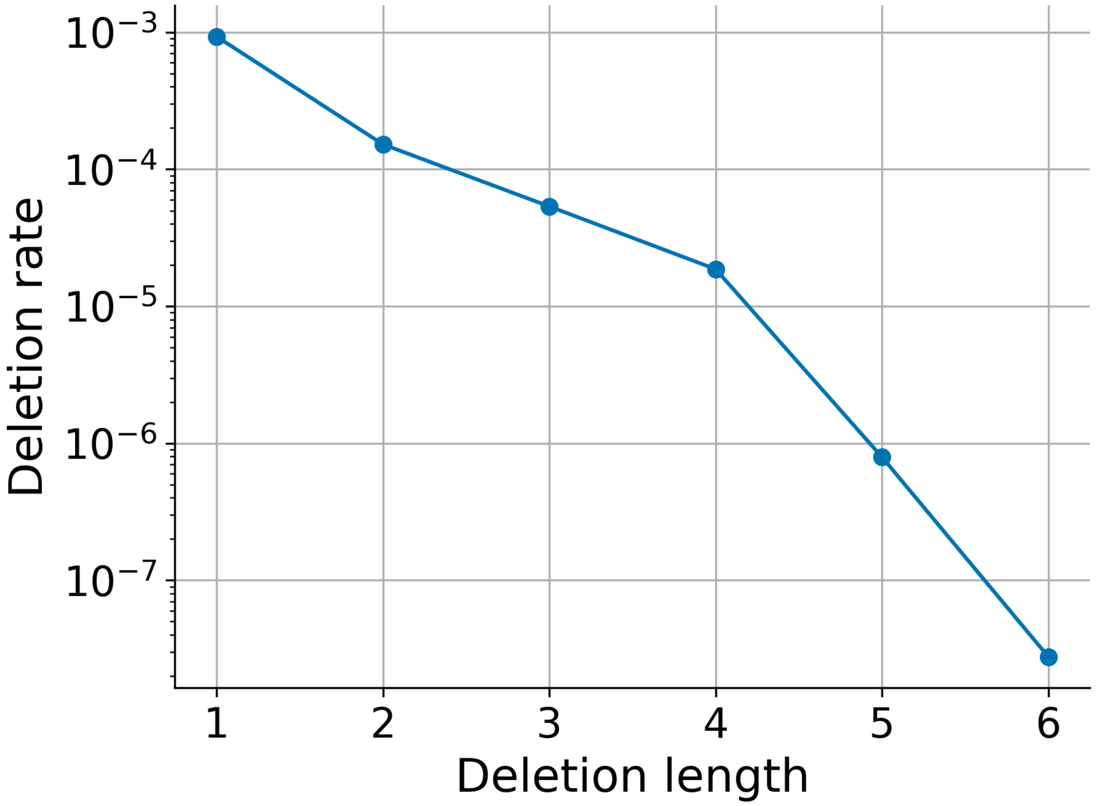

Cummulative distrubtion based upon the number of edit error

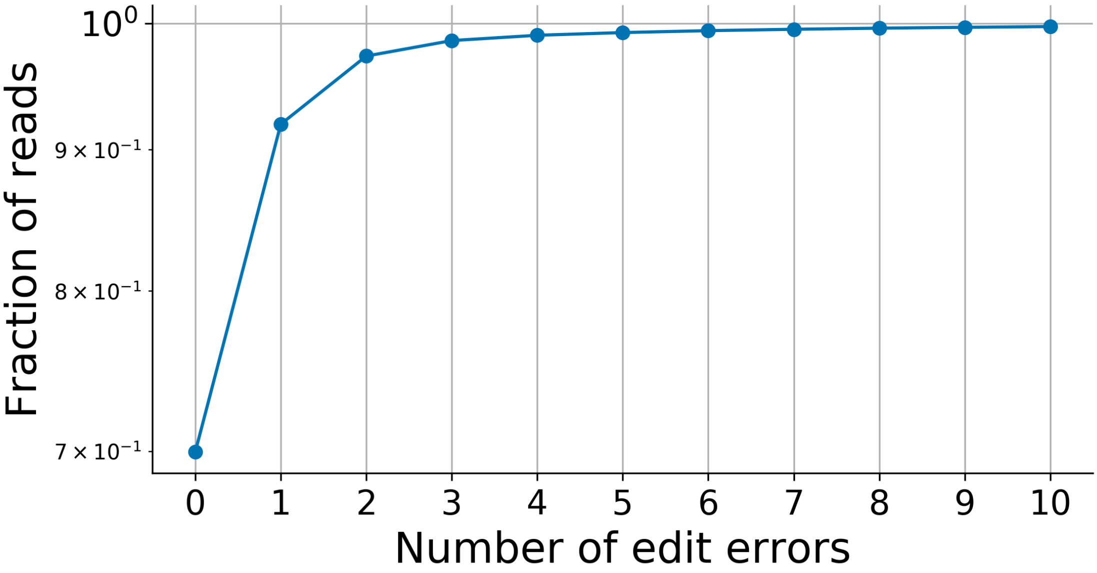

Cummulative distrubtion based upon the number of edit error

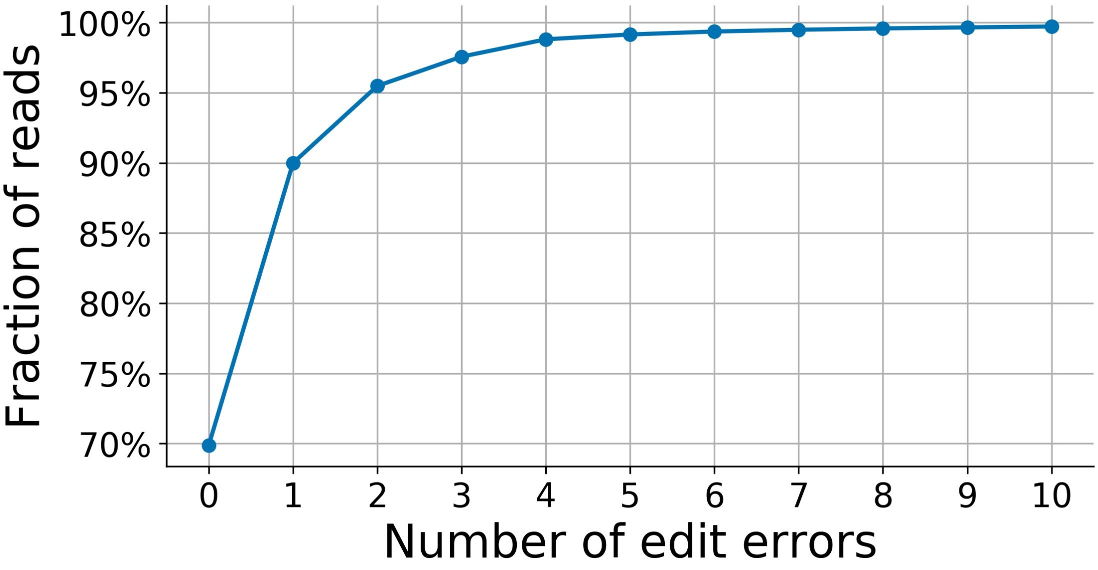

Error rates by GC-Content

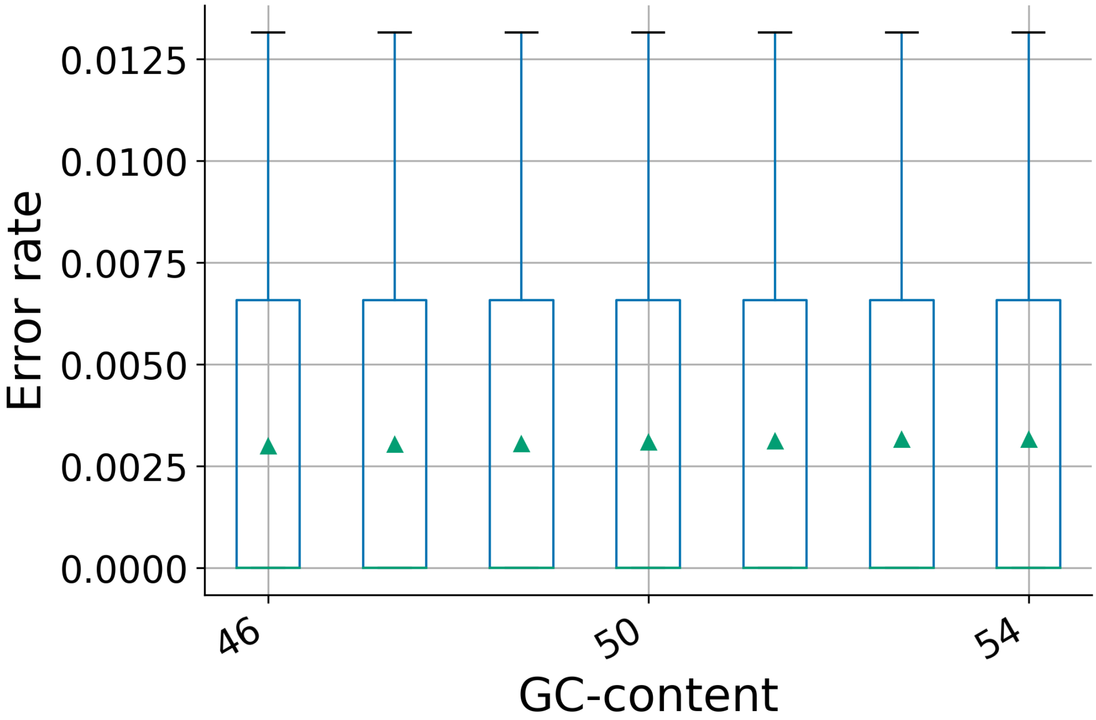
